# How to Improve the Reliability of Aperiodic Parameter Estimates in M/EEG: A Method Comparison

**DOI:** 10.1101/2025.11.10.687541

**Authors:** Patrycja Kałamała, Grace M. Clements, Mate Gyurkovics, Tao Chen, Kathy A. Low, Monica Fabiani, Gabriele Gratton

**Affiliations:** Beckman Institute, University of Illinois Urbana-Champaign, U.S.A.; Psychology Department, University of Illinois Urbana-Champaign, U.S.A.; Centre for Cognitive Science, Jagiellonian University, Poland; Air Force Research Laboratory, Wright-Patterson AFB, U.S.A.; School of Psychology, University of East Anglia, U.K.

**Keywords:** aperiodic activity, 1/*f*, *fooof*, reliability, psychometrics, EEG, censored regression

## Abstract

Interest in broadband aperiodic brain activity (1/*f* phenomenon) has increased exponentially over recent years, partly fueled by the development of tools to parameterize it (i.e., estimate its offset/intercept and exponent/slope) using the M/EEG power spectrum. Broadband aperiodic activity needs to be separated from narrowband periodic activity before its parameters are computed. A popular method, the *fooof* toolbox (Donoghue et al., 2020), is based on the data-driven detection of narrowband-periodic peaks, whose maximum number is set by the user. While increasing analytic flexibility, variability in the number of detected peaks may increase sensitivity to noise and reduce the reliability of aperiodic parameter estimates and the power of analytic pipelines. Here, we present an investigation of the effects of analytic choices (e.g., number of peaks, spectral estimation method) on metrics indicating the adequacy of spectral parametrization. These include the internal consistency (odd-even reliability) of aperiodic estimates, the number of outliers generated, and their ability to detect effects. Across two different data sets (resting state and task-based) we found a decrease in the reliability of intercept and slope estimates as more peaks were allowed to be extracted. To ameliorate this problem, we propose a theory-driven modification of *fooof* labelled *censored regression*, whereby a theory-driven range of frequencies expected to contain periodic activity is removed from all spectra, and the remaining power values are regressed on the remaining frequencies to obtain parameter estimates. This method shows more reliable and robust estimates compared to *fooof*, while avoiding overfitting.

## 1. Introduction

When examined in the frequency domain, the power spectrum of neural activity has a characteristic shape: power decreases as a function of frequency, following a power law of the form 1/*f*^x^. In this equation, *f* represents frequency and *x* is the spectral exponent that determines the rate at which power drops off as a function of frequency (He, 2014; Gao, 2016; Donoghue et al., 2020). This broadband pattern represents non-oscillatory, aperiodic activity, in contrast with neural oscillations, which are narrowband and periodic in nature (Gao, 2016; Donoghue et al., 2020). For decades 1/*f*-like activity was considered “background noise” in electrophysiological signals, with little to no relevance for behavior and cognition. Recently, however, it has been recognized as a signal of interest that changes with age (Voytek et al., 2015; Clements et al., 2021), clinical status (e.g., ADHD; Robertson et al., 2019), sleep patterns (Lendner et al., 2023), and experimental stimulation (Waschke et al., 2021; Gyurkovics et al., 2022; Kałamała et al., 2023). One popular framework links the specific shape of 1/*f* activity to changes in the balance between excitatory and inhibitory inputs within underlying neural circuits (Gao et al., 2017), providing a testable account of and mechanistic interpretation for aperiodic changes (Ahmad et al., 2022).

This newfound interest in aperiodic activity, resulting in a wealth of empirical findings largely in human scalp EEG research, has partly been fueled by methodological developments that allow for the separation and parameterization of broadband 1/*f* activity and periodic, narrowband activity (Wen and Liu, 2016; Donoghue et al., 2020; Kosciessa et al., 2020; Ouyang et al., 2020; Barry and De Blasio, 2021). The *fooof* (fitting oscillations & one-over-f) toolbox (recently renamed *specparam*) in particular has reached immense popularity, as indicated by its over 1500 citations (based on Google Scholar™) within 5 years of its publication (Donoghue et al., 2020). The user-friendly *fooof* algorithm decomposes the power spectrum into periodic and aperiodic parts by first calculating an initial fit of the aperiodic component in log-log space (where both frequency and power are log-transformed), which is then subtracted from the full spectrum (see Gyurkovics et al., 2021, for a discussion of the potential disadvantages of this approach) to create a “flattened spectrum.” Next, peaks exceeding a user-defined noise threshold in the flattened spectrum are fit with Gaussian functions, allowing the determination of each peak’s central frequency, bandwidth, and power corresponding to the center, width, and height of the Gaussian functions, respectively. These fitted Gaussians are then subtracted from the full spectrum, and the final aperiodic fit is calculated on the peak-free remainder using linear regression in log-log space. The slope of this regression captures the steepness of the aperiodic component, while the intercept captures the (base 10) log of the power at 1 Hz. These parameters map onto the “exponent” and “offset” parameters output by *fooof,* respectively. In this article, we use the regression-based terminology (“slope” and “intercept”) as these terms more accurately reflect the estimates used in our calculations, and because their interpretation may be more intuitive to readers less familiar with the aperiodic literature. In all our analyses, we regressed the log of power values onto the log of frequency values (*log10(power) ∼ log10(f)*) as opposed to the log of 1/*f* (*log10(power) ∼ log10(1/f)*). As such, negative slopes indicated a decrease in power with increasing frequency (in line with a typical 1/*f* shape), while positive slopes indicated an increase in power.

Notably, *fooof* requires investigators to choose several parameters, including the threshold used to define each periodic component and the maximum number of possible periodic components (peaks) that can be detected and subtracted (Donoghue et al., 2020). These choices are important because the actual estimates of the broadband aperiodic component may vary depending on how many narrowband periodic components are subtracted. In general, not only will the total power of the aperiodic component be smaller the more periodic components are subtracted, but also the slope of the aperiodic component will vary depending on the number of periodic components subtracted from the lower or higher frequency parts of the spectrum. Especially for high frequencies (where power is expected to be very small), subtracting vs. not subtracting periodic components before estimating the aperiodic parameters may lead to very different results. Further, basing the definition of peaks on particular criteria in the observed spectra (which will likely contain error variance), may also lead to unpredictable outcomes, which may lower the reliability of the 1/*f* parameter estimates.

This problem could be addressed in one of two ways: 1) the Gaussian fitting step may be skipped altogether, which could be accomplished by setting the *fooof* parameter for the number of possible modelled narrowband peaks to *0* or by applying linear regression to the whole log-log spectrum (in this article we labelled these similar procedures “*fooof-0” and “full regression,* respectively); or 2) by excluding sections of the spectra where we may expect narrowband activity to be prevalent on the basis of decades of EEG research and then performing regression on all remaining points (a procedure we here forth label *censored regression*). *Censored regression* ensures there is no difference between spectra regarding which points (i.e., frequency ranges) exactly are identified as containing periodic components, as points with a high probability of periodic activity are excluded (censored) from further analysis for all channels, trials, individuals, etc. While the censored ranges can and should vary from study to study given that the distribution of periodic components may depend, among other things, on sample characteristics (e.g., age), they should always be fixed across individuals *within* a study for the benefits of this method to manifest. Finally, it is worth noting that *fooof* is agnostic to the method used to estimate the input spectra, leaving open the possibility that different spectral decomposition methods could differentially impact the quality of aperiodic parameter estimates, whether they be derived from *fooof* or from *full* or *censored regression*.

The present study thus compares methods for power spectra estimation (i.e., fast Fourier transform, and Welch’s approach for smoothing spectra; referred to as FFT and Welch, respectively), and methods for estimating 1/*f* parameters in log-log space (*fooof*, allowing 0, 1, and 3 narrowband peaks; thereon labeled *fooof0*, *fooof1*, and *fooof3*), as well as the *full* and *censored regression* approaches introduced earlier. We compare the suitability of these different approaches in two data sets, one recorded during rest with open and closed eyes, and another during a multimodal stop-signal task using the following metrics:

a. **Frequency of occurrence of improbable values (outliers)**. Under normal physiological circumstances, the power spectrum is expected to have a characteristic negative going slope, as detailed above (e.g., Gao et al., 2017; Brake et al., 2024). To preview our findings, during exploration of the data we found that in some cases, spectral parametrization methods yielded positive slopes for certain epochs or electrodes. As positive going power spectra are unlikely to have meaningful, brain-based physiological origins in the data sets we analyzed, we explored the nature and origin of such values further, and used their frequency of occurrence as an indicator of the appropriateness of a given method. In other words, these positive values were treated as outliers and all subsequent analyses were carried out twice: with and without positive slopes included.
b. **Reliability (internal consistency)**. Formally, reliability reflects the covariance between repeated measures (e.g., odd and even trials/items) of the same construct (Nebe, Reutter, et al., 2023). Only a few studies have examined the reliability of *fooof* estimates (Levin et al., 2020; Pathania et al., 2021; Lopez et al., 2023; Popov et al., 2023; Tröndle et al., 2023; Webb et al., 2023; Li et al., 2024; McKeown et al., 2024). They mostly focused on test-retest reliability (i.e., *temporal consistency*), with various delays between test and retest, and found moderate-to-good consistency estimates (Lopez et al., 2023). However, *internal consistency* of the estimated parameters, i.e., their reliability within a single measurement period (Cortina, 1993), has rarely been investigated before correlating them across time points (see e.g., Karalunas et al., 2022 for developmental assessment of aperiodic internal consistency). To address this gap, here, we focused on odd-even reliability. Notably, the internal consistency of underlying measures is important because unreliable measures lead to an underestimation of the true correlation, as a substantial portion of the observed interindividual variance is simply due to measurement error. As such, extant results only speak to the psychometric properties of *fooof* estimates indirectly: test-retest correlations may have been moderate either because 1) the measures being correlated had low internal consistency, or 2) because aperiodic parameters are truly not stable trait-like features (or a combination of both). To the best of our knowledge, only one study so far (Karalunas et al., 2022) has addressed the internal consistency of *fooof* parameters using odd-even reliability, and found it to be good-to-excellent in infants and adolescents. Crucially, however, none of these studies (including Karalunas et al., 2022) examined whether reliability changes across *fooof*’s parameter space, i.e., whether there are parameter settings within a reasonable set of potential parameters that consistently yield more vs. less reliable estimates. Consequently, a systematic and in-depth analysis of the reliability of aperiodic *fooof* estimates as a function of various fitting parameters available in the algorithm is missing from the literature. This hinders the research community’s ability to determine best practices (e.g., parameter settings) for using the algorithm, especially in individual differences (correlational) research, where reliability places important constraints on the magnitude of correlations that can be observed with a given measure. Notably, much of the research to date on the role of scalp-level aperiodic activity in cognition has been correlational, such as lifespan or clinical studies (Voytek et al., 2015; Robertson et al., 2019; Clements et al., 2021; McSweeney et al., 2023). An important aspect of reliability is its dependence on the number of trials that are used for parameter estimation: in general, the larger the number of trials, the better the reliability (Hedge et al., 2018; Rouder et al., 2019; Lee et al., 2025). At the same time, increasing the number of trials may engender practical limitations, such as long study duration leading to participants’ fatigue. Therefore, an ideal procedure should produce high reliability estimates (e.g., *r* > 0.9) with a relatively small number of trials/EEG epochs.
c. **Effect size estimates within experimental contexts**. Ultimately, an optimal procedure is one that is most likely to find differences if they truly exist, in other words one that maximizes statistical power. Here, using existing experimental data sets, we opted to capture this by rank ordering effect size estimates recovered by different procedures for effects that have been shown to consistently exist in other, independent data sets. An optimal procedure is one that yields the largest effect size for the same effect. This approach, therefore, rests on the assumption that previously recovered effects are likely to be replicable, and, as such, procedures that can capture them are preferable over procedures that cannot. In the resting state data set, we examined two effect sizes: 1) the magnitude of the correlation between age and the aperiodic slope, where a flattening of the spectrum was expected with increasing age based on several previous studies (Voytek et al., 2015; Dave et al., 2018; Cesnaite et al., 2023; Turner et al., 2023); 2) the difference in slope between the eyes open (EO) and eyes closed (EC) resting conditions, which is an active area of research in the field, with mixed findings (e.g. Euler, Vehar, et al., 2024; Stanyard et al., 2024). While this effect is less crystalized in the literature, we investigated it due to its clear relevance to the discussion around the functional significance of aperiodic activity. We expected the EC condition to be associated with increased inhibition (a steeper spectrum) compared to EO due to the lack of active ongoing visual processing, which requires a cortical configuration ready to receive percepts (see Zhang et al., 2019 for evidence supporting this logic). 3) In the multimodal stop-signal task, we were interested in the effect size of the difference between more and less demanding conditions. Specifically, we expected more demanding conditions to result in steeper spectra (i.e., more negative slopes) compared to less demanding conditions, based on previous findings (Gyurkovics et al. 2022; Kałamała et al., 2024; Lu et al. 2024).

It is important to note that we did not consider the fit of the spectrum to be a key metric of suitability. Fit is the proportion of variance in a spectrum accounted for by the model and is often considered critical to indicate the success of the parameter estimation procedure (e.g., Schaworonkow & Voytek, 2021; Waschke et al., 2021; Gyurkovics et al., 2022; Hill, Clark, et al., 2022; Stanyard et al., 2024). However, fit is expected to increase with the number of free parameters entered in the spectral model. As such, *fooof3* (which includes up to three periodic components alongside the aperiodic component) yielded the highest fit (average spectra median *r*^2^ = 0.96-0.99), while *full* and *censored regression* (which only includes the aperiodic component) resulted in the lowest fit (average spectra *full regression* median *r*^2^ = 0.70-0.86; censored *regression* median *r*^2^ = 0.86 - 0.94). Critically, to the extent that noise variance is also fitted, results may be misleading due to overfitting. Therefore, we considered the above-mentioned alternative metrics (listed in a-c) as our dependent variables of interest and discuss fit only in a limited fashion. Information criterion metrics that combine fit and model complexity can be found in the Supplementary Materials (<u>**Figs. S1** and **S2**)</u>, along with an extended discussion about the difficulties inherent to such metrics when used to adjudicate between methods.

In summary, our study sets out to investigate the effects of common fitting parameter choices and other analytic decisions for the analysis of aperiodic activity, such as the spectral estimation method, on several metrics indicating the adequacy of spectral parametrization. We used two different data sets to this end, one recorded during rest with open and closed eyes, and one during a multimodal stop-signal task. Our findings can help the research community identify critical nodes in their analytic decision tree that could impact the reliability and power of their findings and analyses.

## 2. Methods

### 2.1 Resting State Data set

#### 2.1.1 Participants

Sixty-one adults aged 25-75 participated in the resting state study (mean age ± *SD* = 45.60 ± 15.04, 39 females). All participants were native English speakers and reported themselves to be in good health with normal hearing and normal or corrected-to-normal vision, and free from medications that may directly affect the central nervous system. All participants signed informed consent and all procedures were approved by the University of Illinois Urbana-Champaign Institutional Review Board. Nine participants had excessively noisy data in their reference electrodes, which permeated all other channels, rendering their data inadequate for EEG analysis and were thus excluded from analysis leaving a final sample size of N=52.

#### 2.1.2 Resting State Procedure

Participants sat in a dimly lit, sound- and electrically attenuated recording chamber, and were instructed to sit quietly and not think about anything in particular. Each session included a 5-minute recording of resting EEG with eyes-open followed by 1-minute with eyes-closed. During the eyes open condition, participants fixated on a cross presented centrally on a computer screen. These data were recorded at the beginning of a session that involved other experiments not reported here.

#### 2.1.3 EEG Data Acquisition and Preprocessing

EEG and EOG were recorded continuously from 64 active electrodes mounted in an elastic cap (Acti-Cap) using a BrainAmp recording system (BrainVision Products GmbH) and were filtered online using a 0.5 - 250 Hz bandpass. The sampling rate was 500 Hz. Data were referenced offline to the average of the left and right mastoid electrodes. Preprocessing began by shortening the eyes-open data to 1-minute in length – the first minute of the recording – to be directly comparable to the eyes-closed data. This was followed by the application of a 40-Hz low-pass filter, segmentation of the 1-minute contiguous data into 1024-ms long epochs to facilitate the use of our artifact detection scripts, and ocular artifact correction using the method proposed by Gratton et al. (1983). After eye movement correction, epochs with voltage fluctuations exceeding 200 μV were excluded from further analysis.

### 2.2 Stop-Signal Data set

#### 2.2.1 Participants

The stop-signal task data were collected as part of a separate project conducted at Jagiellonian University in Krakow, Poland, and were made available for the present analysis. Thirty-four young adults participated in the study (mean age ± *SD* = 21.06 ± 1.48, 28 females). All participants were native Polish speakers who reported being in good health, with normal hearing and normal or corrected-to-normal vision, and no use of medications known to directly affect the central nervous system. Written informed consent was obtained from all participants, and all procedures were approved by the Jagiellonian University Review Board.

#### 2.2.2 Experimental Task and Procedure

Participants completed four versions of the stop-signal task, which differed in the modality of both go and stop stimuli, yielding the following variants (go-stop, respectively): (1) visual-visual, (2) auditory-auditory, (3) visual-auditory, and (4) auditory-visual.

Each trial began with the presentation of a white central fixation cross for 1200 ms, immediately followed by the presentation of a blank screen for 200 ms. After, the go stimulus was presented for 100 ms. In a random sample of 25% of the trials, the stop-signal was presented, prompting participants to inhibit their responses to the primary go task. The interval between the presentation of the go stimulus and the stop-signal (stop-signal delay, SSD) was varied trial-by-trial using a tracking method: the interval increased or decreased by 50 ms (from 100 to 400 ms) for the subsequent stop-signal trial, depending on whether the participants successfully or unsuccessfully inhibited their response to the go stimulus. Thus, there were seven possible SSDs: 100, 150, 200, 250, 300, 350, 400 ms. The initial SSD value was 150 ms. The tracking method aimed to converge on an SSD at which participants successfully inhibited responses in approximately 50% of the stop-signal trials. The timeout for a trial was 1500 ms, and the inter-trial interval was 1100 ms. All stimuli were presented against a black background, with participants seated approximately 80 cm from the computer screen. Auditory stimuli were delivered through Sennheiser HD 429 headphones (intensity 60 dB SPL, duration 100 ms, rise and fall time 10 ms).

The *go* stimulus was always presented laterally. The *visual go* stimulus consisted of a green square randomly appearing on the left or right side of the screen, with its inner edge positioned 2° from the center of the screen. The *auditory go* stimulus was a 1000 Hz sound randomly presented to the left or right ear. Participants were instructed to indicate the location of the *go* stimulus (left vs. right side for visual go and left vs. right ear for auditory go) by pressing the left or right “CTRL” key using their index fingers. The *stop* stimulus was always presented bilaterally. The *visual stop* stimulus consisted of two small red squares simultaneously presented on both sides of the screen, with each square’s inner edge positioned at 2° from the center of the screen. The *auditory stop* stimulus involved a 1400 Hz sound presented simultaneously to both ears.

Task order was pseudorandomized. Each task version comprised four experimental blocks, each with 50 trials, resulting in 200 trials per version (800 trials in total). In addition, participants completed one or two practice blocks, each consisting of 24 trials, before engaging in the main task. The task was implemented using *DMDX* software (Forster & Forster, 2003).

#### 2.2.3 EEG Data Acquisition and Preprocessing

EEG and EOG were recorded continuously from 38 active electrodes mounted in an elastic cap using a BioSemi ActiveTwo recording system, at a sampling rate of 256 Hz. Data were referenced offline to the average of the left and right mastoids and filtered with a 0.5-40-Hz band-pass filter. They were then segmented into 1000-ms long epochs (i.e., the final 1000 ms of the fixation cross presentation preceding the stimulus onset). Ocular artifact correction was implemented through the method described in Gratton et al. (1983). After eye movement correction, epochs with voltage fluctuations exceeding 80 μV were excluded from further analysis. For the present study, both correct and incorrect trials were retained for analysis (average number of artefact free epochs across participants = 195, *SD* = 7, *min* = 165, *max* = 200).

### 2.3. Statistical Analyses

Both resting state and stop-signal data sets underwent identical spectral decomposition and aperiodic parameterization procedures, described below. **Table 1** summarizes the technical parameters of signal recording for both data sets.

**Table 1.**
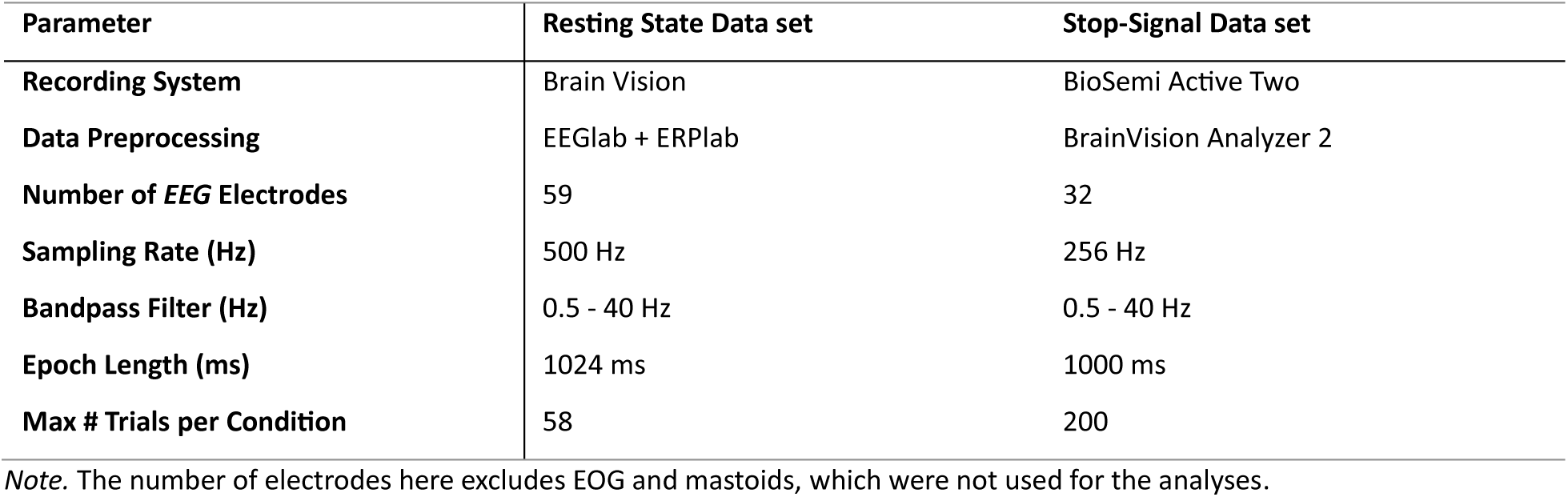
Summary of the technical parameters of the analyzed data sets.

#### 2.3.1. Quantifying the power spectra

We used two common spectral decomposition methods to generate power spectra: the Fast Fourier Transform (*FFT*) and Welch’s method (*Welch*). Spectra derived via *Welch* used the MATLAB default settings. Each method was applied to the epoched data at the single trial level (resulting in a spectrum per trial per electrode per participant) and at the average level (resulting in a spectrum per electrode per participant) using MATLAB’s *fft()* or *pwelch()* functions. The power values from *FFT* and *Welch* were computed up to the Nyquist frequency, but the spectra were subsequently shortened to a range of 2-33 Hz for analysis. The lower bound of the frequency range was determined by the length of our epochs: with ∼1 second of data, a 2 Hz waveform is the slowest that can still resolve multiple (i.e., 2) cycles. The upper bound of the frequency range was determined by the dynamic range of the A:D converter^1^, which in our case was 0.1 µV/bit, indicating that a waveform smaller than this value is indistinguishable from noise in the EEG signal. We determined that 33 Hz was the frequency at which we expect power to be twice as large as our amplifier can detect (0.04 µV ^2^). In other words, at 33 Hz we expect the EEG signal to have an amplitude of 0.2 µV, which is double the dynamic range of our amplifier.

#### 2.3.2. Spectral parameterization

All spectra were parameterized to determine the aperiodic parameters, slope and intercept, using a data-driven approach (i.e., *fooof*) and a theory-driven approach (*full* or *censored regression*). *Fooof* models were fit with 0, 1, or 3 peaks and with the following parameters: *aperiodic_mode* = ‘fixed’, *peak_width_limits* = [2.0, 12.0]. The *fooof0* model served as a baseline to assess the outcome when *fooof* did not remove any periodic peaks from the data. In total, 12 possible *fooof* models were fit: *Data Type (Single, Average) × Spectral Decomposition (FFT, Welch) × Number of Peaks Modelled (0, 1, 3)*, where “single” and “average” mean whether we are investigating the average of single-trial aperiodic estimates or the aperiodic estimates of the average of single-trial spectra, respectively. As such, for single-trial data, spectra were parameterized first, then the estimates were averaged. For average data, spectra were averaged first, then parameterized.

A theory-driven regression approach was also used to parameterize the spectra (a version of which was first used in Clements et al., 2021). Two regression models were used: *full* and *censored*. Both methods follow the same principal steps, spectra were first transformed from linear space into log-log space, and then a single, simple linear regression was fit to the data using MATLAB’s *fitlm()* function. *Full regression* models fit a simple linear regression to the entire “full” power spectrum. In this way, *full regression* served as a ‘baseline’ similar to the *fooof0* model to quantify the slope and intercept naively, using all available data. As noted in the Introduction, *censored regression* is theoretically informed by the myriad extant data indicating that most scalp-recorded periodic activity during wakefulness occurs in the theta-alpha-beta frequency range (e.g., Myrov et al., 2024). Thus, in *censored regression,* power values in the 6–16 Hz frequency range were excluded in the regression model and then the slope and intercept were estimated. This approach excludes frequencies most likely to contain periodic activity during fitting, yielding slope and intercept estimates that are unaffected by such activity. Crucially, the fact that it is always the same frequency range that is omitted (i.e., on each epoch and/or for each electrode) guarantees that the same number of (unadjusted) points are always used to determine aperiodic parameters. See **Fig. S3** in the supplementary materials for a demonstration that this approach yields accurate aperiodic parameter estimates even if the omitted (censored) range contains points uncontaminated by oscillatory peaks, as long as the beginning and the end of the fitting range are left uncensored. In sum, 8 possible regression models were fit: *Data Type (Single, Average) × Spectral Decomposition (FFT, Welch) × Regression Type (Censored, Full)*.

In the *censored regression* approach, log power values were regressed onto the log of frequency values (rather than the log of *1/f*). Because our predictor was *f* and not *1/f*, slope values were expected to be negative, indicating negative-going spectra. Exponent values provided by *fooof* were multiplied by -1 to account for the fact that the algorithm outputs slopes for log(1/*f*), not log(*f*), making the *fooof* exponents into slopes and thus comparable across estimation methods.

The data were analyzed and visualized in *R* 4.0.3 (R Core Team, 2021). The materials, data, and *R* code for this project will be openly available in the project repository https://osf.io/z3t5m/ (https://osf.io/z3t5m/). The analyses described below took into consideration the following factors: Task Type (eyes open and eyes closed for the resting-state data set and visual-visual, visual-auditory, auditory-visual, auditory-auditory for the event-related data set); Spectral Decomposition (*FFT*, *Welch*); Aperiodic Estimation (*fooof0*, *fooof1*, *fooof3*, *regression-censored*, *regression-full*); Channel (59 unique EEG channels in the resting state and 32 in the event-related data set). Statistical tests were performed on all possible pairwise combinations of aperiodic estimation methods using two-tailed tests. In the bar plots, the methods were always ranked from best to worst performing for a given outcome measure, with bracket connections indicating statistically different methods. For all analyses, *p*-values < .05 (without correction for multiple comparisons) were considered significant; this was considered appropriate because the focus here is on identifying optimal methodologies rather than on making theoretical inferences. Next, we outline how each outcome measure (listed as points a-c in the Introduction) was evaluated.

#### 2.3.3. Frequency of improbable values (outliers)

First, we compared the aperiodic estimation methods in terms of their susceptibility to generate positive slope values. As described in the Introduction, during our initial exploration of the data, we identified some data segments within individuals where slope values were positive. These values are neurophysiologically implausible and highly unlikely to occur under normal circumstances. Therefore, we interpreted these outcomes as due to measurement error. First, we quantified the number of epochs with positive values for each combination of *Task Type × Spectral Decomposition × Number of Peaks/Regression Type* for each channel and participant separately. Epoch counts were then averaged across channels for each participant and presented as a percentage relative to the total number of retained epochs. Two non-parametric tests were conducted to compare aperiodic estimation methods in their tendency to produce positive slopes for each task type separately. First, we applied a permutation test with 1000 permutations. We calculated the difference between percentages across participants between a given method pair. Then on each iteration, the sign of the difference values was reversed for a random half of the data points and averaged to build a null distribution. Then, the observed average difference values were compared to the null distribution. Parameter estimates were deemed significantly different between pairs of levels if the observed difference exceeded the top 97.5% or fell below the bottom 2.5% of the null distribution, corresponding to *p* < .05 (two-tailed). Second, we applied a nonparametric bootstrap procedure on the same data. Participants’ data were resampled with replacement 5,000 times. In each bootstrap sample, we calculated the mean percentage of positive slopes per aperiodic estimation method, and ranked methods such that lower values were better (reflecting fewer outlier estimates). We then summarized (1) the probability of being best (lowest % positive slopes) and (2) the consensus mean rank across iterations. This analysis provides a robust, rank-based assessment of which methods are least prone to producing spurious or physiologically implausible outcomes.

We also explored whether positive slope values reflected power spectra that were truly positive going (e.g., because of muscle noise contamination at higher frequencies) or if they were merely artifacts of the analytic procedures used. To this end, we correlated each power spectrum with a negative going line in log-log space across the frequency range of interest (a “mirrored” identity line). Power spectra for which this correlation was negative were taken to be truly positive going. Next, we investigated the proportion of positive slopes accounted for by truly positive spectra for each method. Any unaccounted-for positive slope values were taken to reflect undesired artifacts of the aperiodic estimation procedure. The proportion of such slope values was then used as an additional metric of aperiodic estimation performance, with lower proportions reflecting better performance.

For all subsequent analyses (2.3.4-2.3.5) the impact of positive slope removal was also investigated to highlight the potential importance of this preprocessing step. Whenever results substantially differed from or directly contradicted the main findings, i.e., those before removing positive slope values, the filtered results are also presented alongside the full results. For all other cases these results can be found in the <u>Supplementary Materials</u>. Notably, for the filtered data, all positive slopes were removed, not just those unaccounted for by the presence of positive going spectra.

#### 2.3.4 Reliability (internal consistency)

Single-trial data served to assess the psychometric properties of the aperiodic component. In the first step, the internal consistency of the slope and intercept were estimated as the Spearman’s rho correlation (due to the skewed distribution of the data) between odd and even numbered epochs. Again, two non-parametric tests were applied. A permutation test (described in 2.3.3) was used to test significant differences in reliability among the aperiodic estimation methods for each Task Type separately. A bootstrap test (similar framework described in 2.3.3) for each task condition (version in the stop-signal data set; eyes open/closed in the resting state data) was applied. Participants’ data were resampled with replacement 5,000 times. Within each bootstrap iteration, we computed the mean reliability coefficient (odd–even correlation, ρ) for each aperiodic estimation method and established a rank order (1 = best, higher = better). For every method, we recorded (1) the probability of being the best (percentage of bootstrap samples in which it ranked 1st), and (2) the consensus mean rank (average rank across all iterations). This bootstrap ranking approach quantifies both the likelihood that a method provides the most consistent estimates, and the stability of its relative position among alternatives.

In addition to the conventional odd-even reliability, we also investigated the minimal number of epochs required to obtain reliability estimates greater than .90 for the slope and intercept. For this purpose, we computed the odd-even *rho* coefficients using the aforementioned procedure, varying the number of epochs incrementally, starting from the first ten epochs in the data, and increasing by increments of ten (6 iterations for the resting-state data set and 11 iterations for the event-related data set).

#### 2.3.5. Effect size estimates within experimental contexts

In addition to their reliability, we also examined which of the aperiodic estimation methods was most effective in detecting the effects of interest, i.e., which one was most likely to be useful for the research community. As noted in the Introduction, these were 1) the relationship between age and slope in the resting state data, 2) the difference between EO and EC in the resting state data, and 3) the effect of cognitive demand on the steepness of the spectrum in the task-based data. For 1, effect size was captured by Spearman’s *rho*. For 2 and 3, the effect size measure was Cohen’s *d*. For these analyses, single-trial aperiodic estimates were averaged across epochs and then across channels. In general, methods that resulted in larger effect sizes in the hypothesized directions were considered to be more valid than those resulting in smaller effect sizes.

## 3. Results

### 3.1. Descriptive statistics

**Table 2** includes descriptive statistics for both data sets. The results show that the different methods produce slightly different mean slope and intercept values. It should be noted that the maximum slope values for some analytic approaches are positive, which is a concern that will be addressed later in section 3.2. *Table S1 p*resented in the <u>Supplementary Materials</u> contains the same descriptive information after the removal of positive slope values.

**Table 2.**
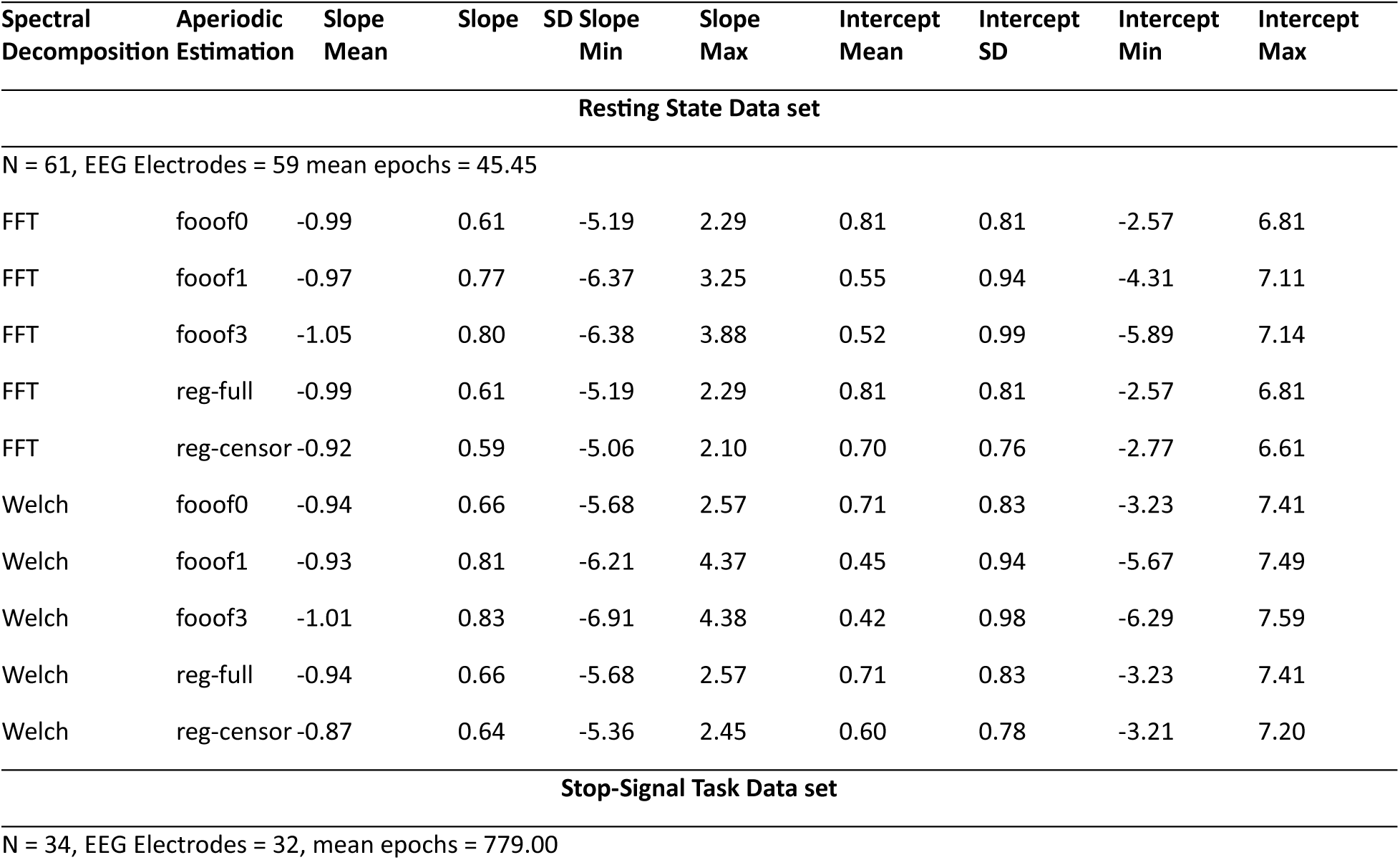
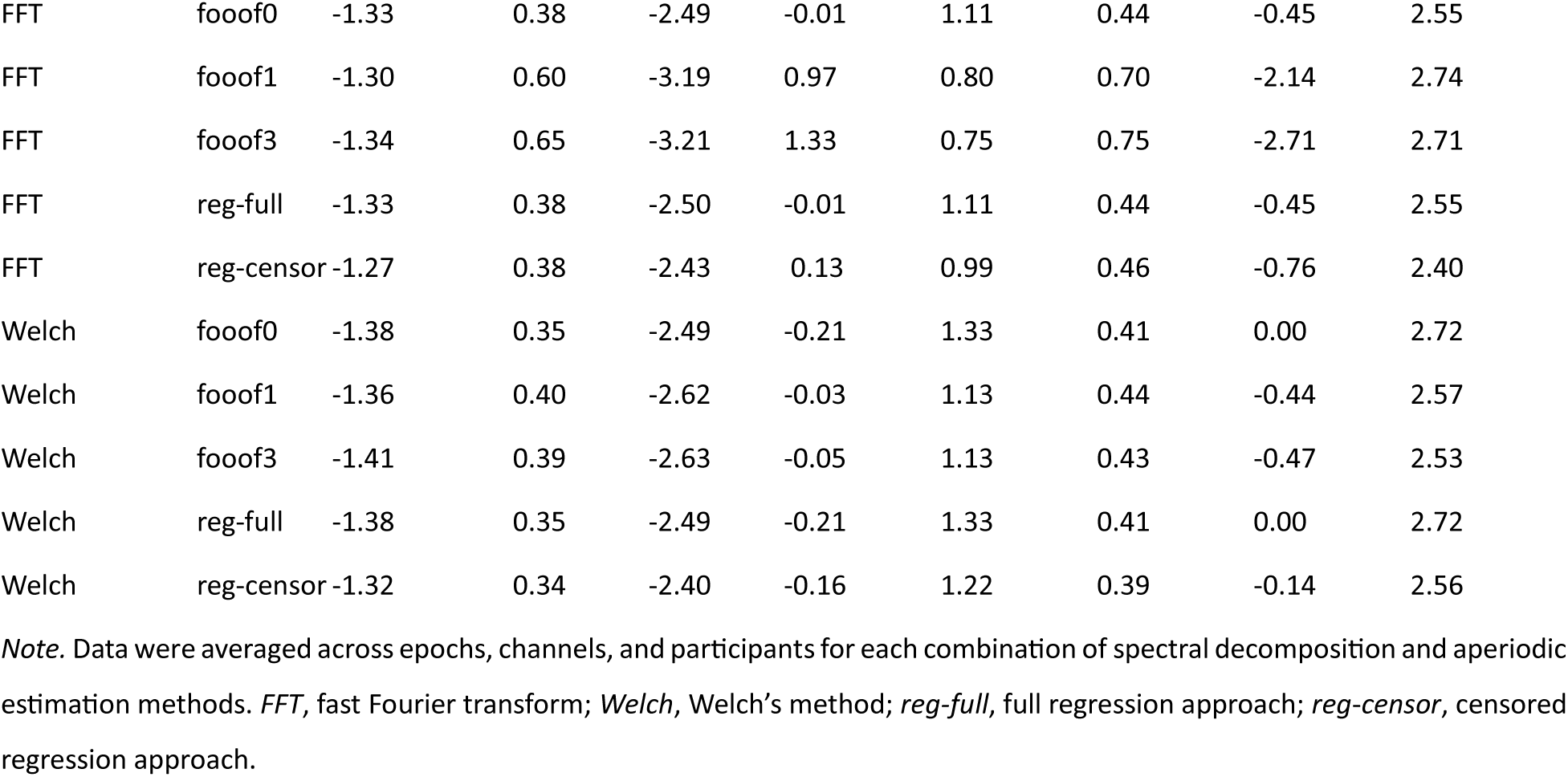
Average slope and intercept values across participants and electrodes for each estimation approach.

Complementing the descriptive data, the total power spectra and the reconstructed aperiodic components in log-log space for the resting state data and one version of the Stop-Signal Task (visual *go* followed by visual *stop*) are shown below in **Fig. 1** at the midline electrodes Fz, Cz and Pz. Using FFT or Welch primarily impacts the total power spectrum at the lowest frequencies, albeit only marginally so (**Fig. 1A**). For most analytic approaches, aperiodic reconstructed power spectra are highly comparable between single trial data and average data, with only some approaches producing slightly different (typically higher) intercepts (broadband shifts) for the averaged compared to the single trial data set. This is particularly evident in the *fooof*-derived resting state components with one or more peaks (see the first six columns of the top two panels of Fig. 1B). These figures suggest that different analytic decisions produce generally convergent spectra under the settings considered here, with only relatively small differences in either the full spectrum or the aperiodic component only.

**Figure 1.**
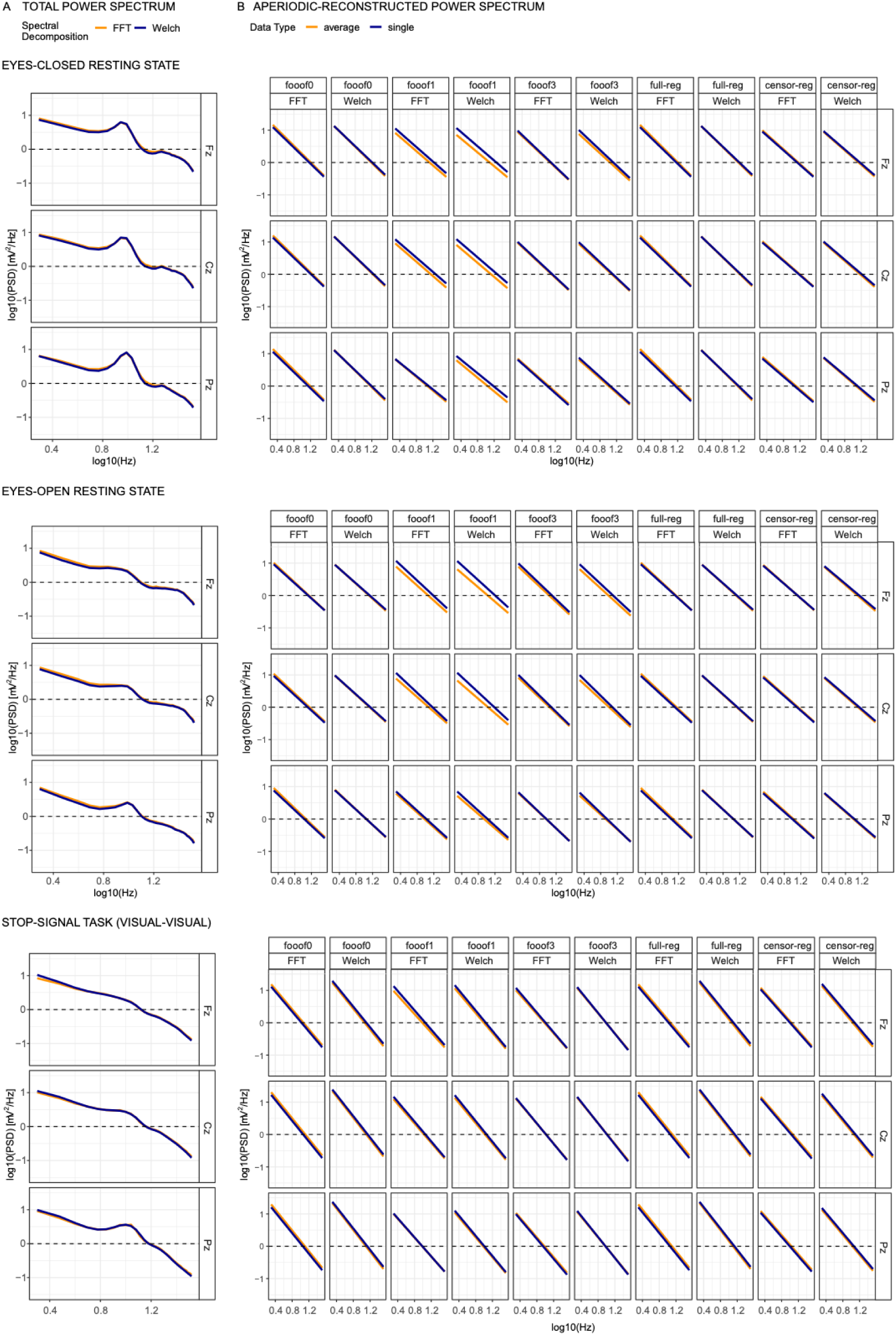
Actual (A) and modelled (B) power spectra in log-log space for the eyes-closed resting state (top), eyes-open resting state (middle), and the visual-visual version of the stop-signal task (bottom), shown across three selected midline electrodes: frontal (Fz), central (Cz), and parietal (Pz). Column B presents all combinations of spectral decomposition and aperiodic estimation methods. Only the aperiodic component was modelled for each cell in B. *FFT*, fast Fourier transform; *Welch*, Welch’s method; f*ooof0*, *fooof1*, and *fooof3*, *fooof*-derived slope estimates with the corresponding number of modelled peaks; *full-reg*, full regression approach; *censor-reg*, censored regression approach.

### 3.2 Frequency of occurrence of improbable values (outliers)

After comparing descriptive statistics across pipelines, we turn to the question of improbable values – positive slopes – generated by each method. The percentage of epochs with positive slopes produced by the various methods in the resting state and stop-signal task data sets are shown in **Fig. 2**. In both data sets, *fooof1* and *fooof3* led to the highest proportion of positive slopes. This pattern was observed regardless of spectral decomposition method in the resting state data (**Fig. 2**, top two panels, left columns), whereas it only emerged for *FFT* in the stop-signal data set. Notably, after accounting for genuine positive going spectra (see 2.3.3), epochs with positive slopes clearly diminished for all methods, except *fooof1* and *fooof3* (right columns of **Fig. 2**), suggesting an occasional misestimation of real slope values by these methods. A few outliers still remained for all but the *censored regression* method. Permutation testing and bootstrap analyses generated almost identical results. Therefore, for simplicity, results described here and in **Fig. 2** reflect permutation testing. Bootstrap analyses can be found in the <u>Supplementary Materials,</u> ***<u>Tables S2</u>* and *<u>S3</u>***.

**Figure 2.**
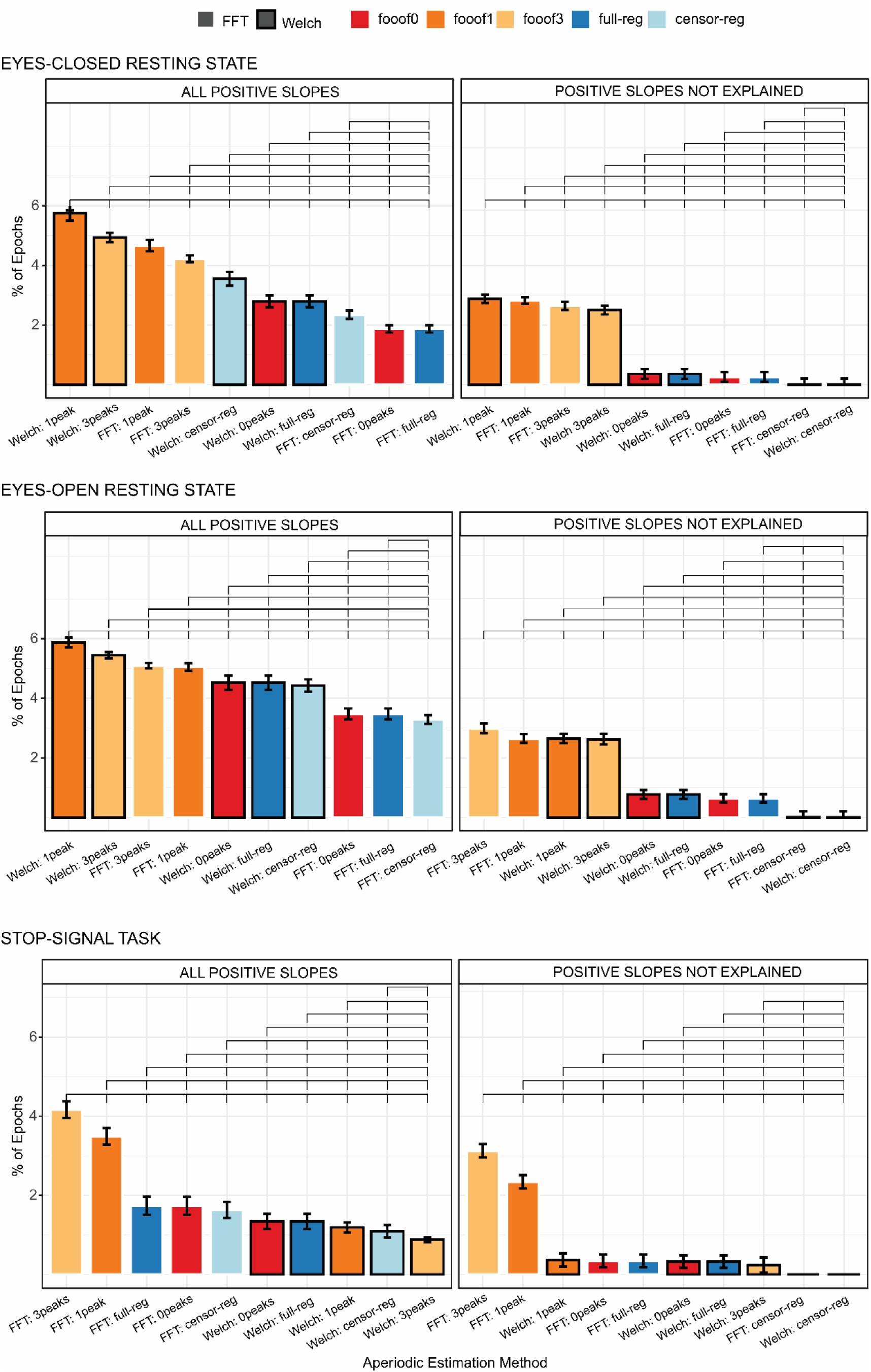
Mean percentage of positive slopes before (left column) and after removing genuinely positive going spectra (right column), for all combinations of spectral decomposition and aperiodic estimation methods for the eyes-closed (top panel), eyes-open (middle panel), and stop-signal (bottom panel) conditions. Within each subplot, bars are ordered from highest to lowest value. Brackets above the bars indicate statistically significant differences (*p* < .05 using permutation testing) and error bars reflect within-subject standard errors of the mean (SEM), computed using Morey’s (2008) correction for repeated-measures designs. *FFT*, fast Fourier transform; *Welch*, Welch’s method; *fooof0*, *fooof1*, and *fooof3*, *fooof*-derived slope estimates with the corresponding number of modelled peaks; *full-reg*, full regression approach; *censor-reg*, censored regression approach.

### 3.3 Reliability (internal consistency)

#### 3.3.1 Resting State

Odd-even reliability derived from the single-trial resting state data before outlier removal is presented in **Fig. 3** for slope and intercept and each eye status condition (closed/open). For slope, the analytic approaches that produce the highest reliability are the *FFT*-based data with *full regression* and *fooof0* (two methods that are practically identical to each other and do not distinguish between periodic and aperiodic activity), regardless of eye status. Internal consistency is still greater than 0.9 for *FFT*- or *Welch*-based data using *fooof0*, *full regression* and *censored regression*. Including more peaks in the *fooof* models significantly decreases the odd-even reliability for the slope. Similarly, for the intercept, *FFT*-based data with *full regression* or *fooof0* produce the highest reliability, followed by *Welch*-based data, and then *censored regression* with either *FFT* or *Welch*. Allowing for more peaks to be included in the *fooof* models significantly reduces internal consistency across both parameters (0.86 or less). Results after the removal of positive slopes were entirely consistent with these conclusions, producing numerically similar results, suggesting that the likelihood of estimating a positive slope is also highly reliable. These results are shown in ***Fig. S4*** in the <u>Supplementary Materials</u>. As in section 3.2., permutation testing and bootstrap analyses generated almost identical results. Therefore, for simplicity, results described here and in **Fig. 3** reflect permutation testing. Bootstrap analyses can be found in the <u>Supplementary Materials</u> ***Tables S4-S5***.

**Figure 3.**
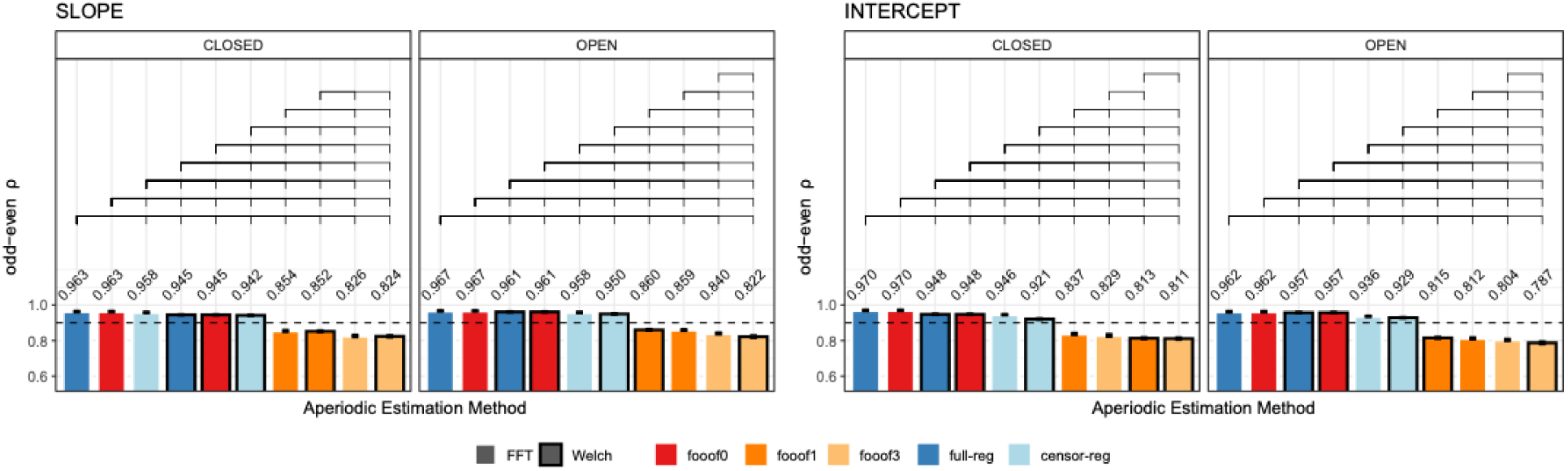
Internal consistency (odd-even reliability) for slope (left) and intercept (right) across aperiodic estimation methods, shown separately for eyes-closed and eyes-open resting state. Within each subplot, bars are ordered from highest to lowest reliability. Brackets above the bars indicate statistically significant differences (*p* < .05, permutation testing) and error bars reflect within-subject standard errors of the mean (SEM), computed using Morey’s (2008) correction for repeated-measures designs. The horizontal dashed line represents a reliability threshold of .90, considered acceptable. *FFT*, fast Fourier transform; *Welch*, Welch’s method; *fooof0, fooof1*, and *fooof3*, *fooof*-derived slope, *full-reg*, full regression approach; *censor-reg*, censored regression approach.

Next, we assessed the number of trials needed to obtain reliable estimates of slope and intercept (**Fig. 4**). The data from *full regression* and *fooof0* are overlapping, as expected. Importantly, fewer trials are needed to obtain reliable estimates for the non-*fooof* regression methods (blue lines) compared to *fooof* with any peaks (orange-red lines), as well as fewer for the slope compared to the intercept. To obtain slope estimates with > 90% reliability using non-*fooof* regression methods, ∼30-40 trials are needed, whereas >90% reliability for the intercept required all 60 trials in the resting state data sets. More than 60 trials (∼1 min of recording) would be required for either parameter to reach excellent reliability using *fooof* with 1 or more peaks. Results after positive slope removal are fully in line with these conclusions and can be found in ***Fig. S5*** in the <u>Supplementary Materials</u>.

**Figure 4.**
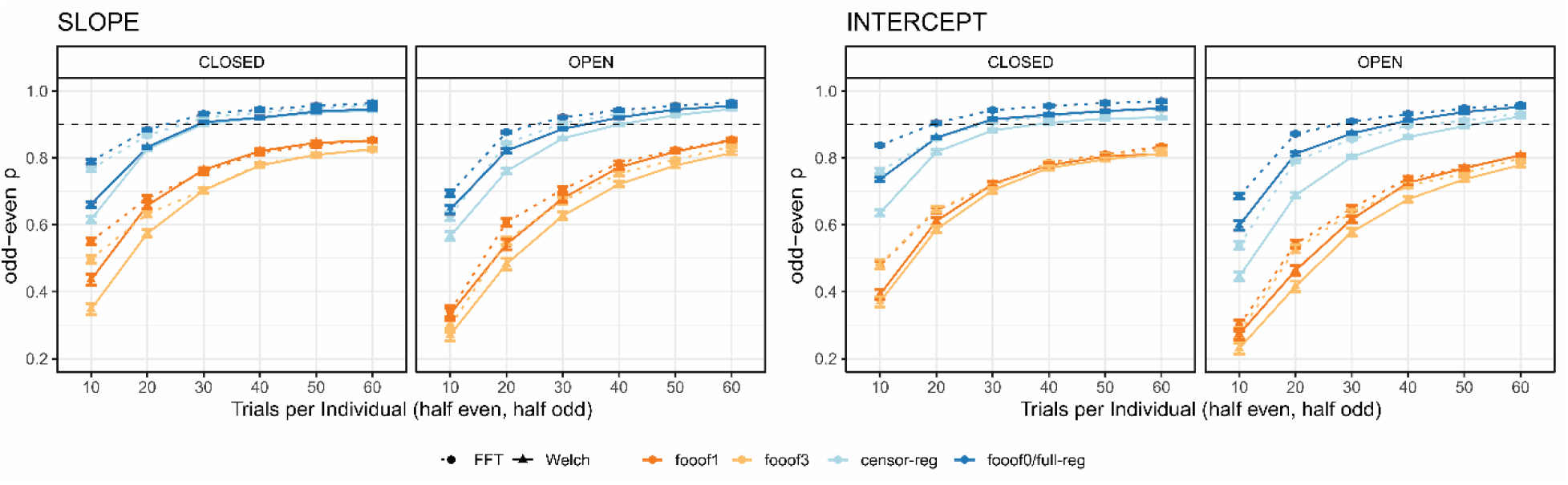
Internal consistency (odd-even reliability) as a function of trial count for slope (left) and intercept (right) across aperiodic estimation methods, shown separately for eyes-closed and eyes-open resting state. The dashed horizontal line represents a reliability threshold of .90, considered acceptable. fooof0 and full-reg are displayed with the same color because their curves were virtually indistinguishable from each other. FFT, fast Fourier transform; Welch, Welch’s method; fooof0, fooof1, and fooof3, fooof-derived slope estimates with the corresponding number of modelled peaks; *full-reg*, full regression approach; *censor-reg*, censored regression approach.

#### 3.3.2 Stop-Signal Task

Odd-even reliability derived from the single-trial data are presented in **Fig. 5** for slope and intercept estimates and each trial type in the stop-signal task. For the slope, the analytic approaches that produce the highest reliability are consistently *Welch*-based data with *full regression*, then *fooof0*, then *censored regression*, followed by *FFT*-based data with *full regression*. Across all conditions, *fooof* models with 1 or more peaks consistently ranked lowest. Reliability for the intercept estimates showed comparable, albeit less consistent patterns, where *fooof0*, *full regression* and *censored regression* tended to be ranked significantly higher than any *fooof* models with non-zero peaks. Results after positive slope removal are in line with the general conclusion that multi-peak models perform worse in all conditions, although the exact ranking of specific top methods shows some variability (*Fig. S6*, <u>Supplementary Materials</u>). As before, permutation testing and bootstrap analyses generated almost identical results. Therefore, for simplicity, results described here and in **Fig. 5** reflect permutation testing. Bootstrap analyses may be found in the <u>Supplementary Materials, *Tables S6-S9*</u>.

**Figure 5.**
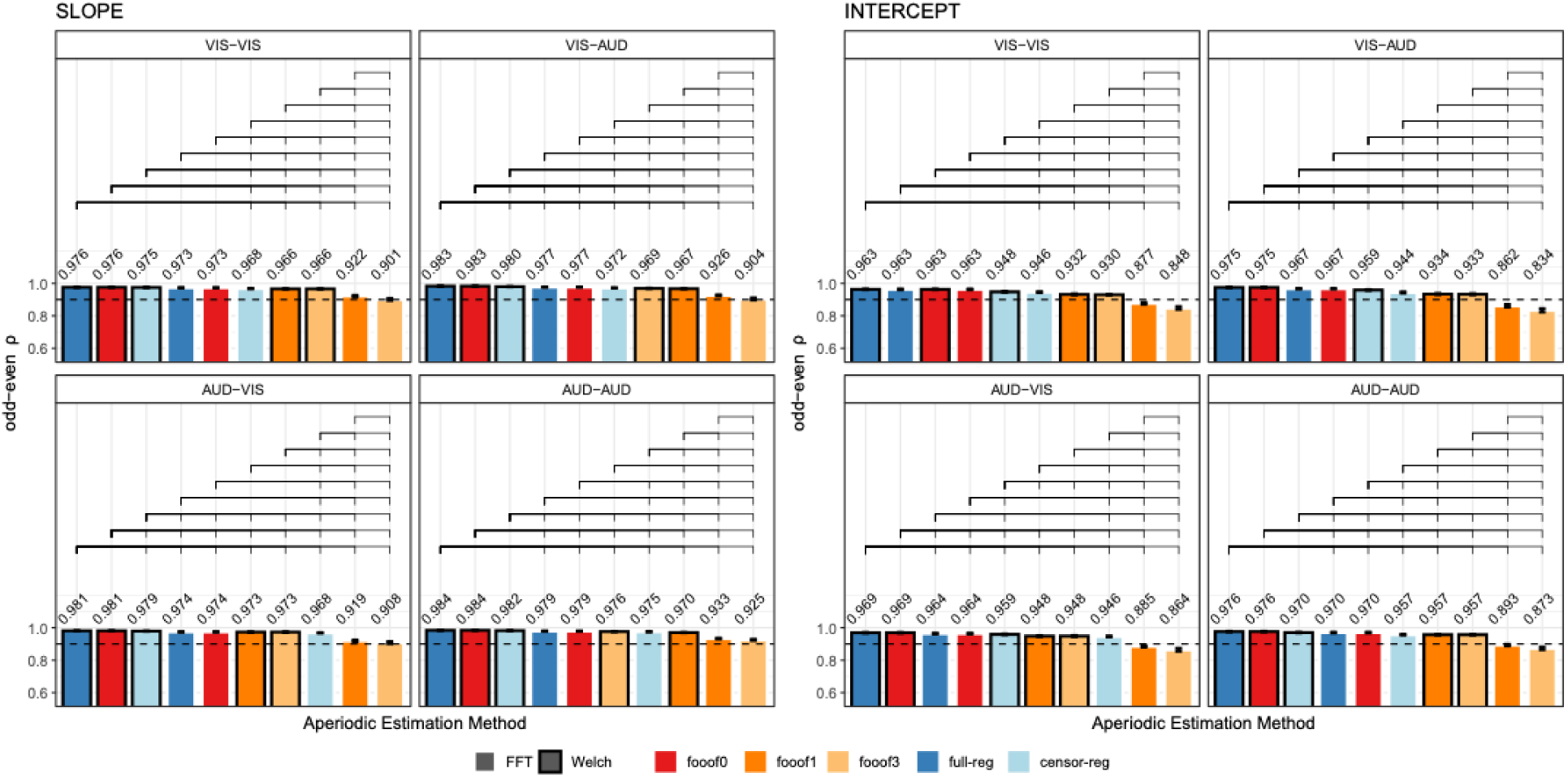
Internal consistency (odd-even reliability) for slope (left) and intercept (right) across aperiodic estimation methods, shown separately for each version of the stop-signal task. Within each subplot, bars are ordered from highest to lowest reliability. Brackets above the bars indicate statistically significant differences (*p* < .05 permutation testing; see text for details). Error bars reflect within-subject standard errors of the mean (SEM), computed using Morey’s (2008) correction for repeated-measures designs. The horizontal dashed line represents a reliability threshold of .90, considered acceptable. *FFT*, fast Fourier transform; *Welch*, Welch’s method; *fooof0*, *fooof1*, and *fooof3*, *fooof*-derived slope estimates with the corresponding number of modelled peaks; *full-reg*, full regression approach; *censor-reg*, censored regression approach.

Next, we assessed the number of trials needed to produce reliable estimates of the aperiodic parameters (**Fig. 6**). Across all four conditions and for both slope and intercept, the general conclusion from the resting state data holds, both before and after positive slope removal (***Fig. S7***) – fewer trials are required to reach .90 reliability for the *censored* and *full regression* approaches and *fooof0* compared to *fooof1* and *fooof3*. However, in this data set, *Welch*-based methods clearly outperformed *FFT* at least for *fooof* models with non-zero peaks: while slope estimates in all other pipelines reached excellent reliability with well less than 100 trials, *fooof1* and *fooof3* with *FFT* often needed as many as 130 trials to cross this threshold. For the intercept parameter, all pipelines required more trials than the slope to achieve high reliability, but the inferiority of *FFT*-based approaches remained similar, in some cases requiring more than 190 trials to reach the threshold.

**Figure 6.**
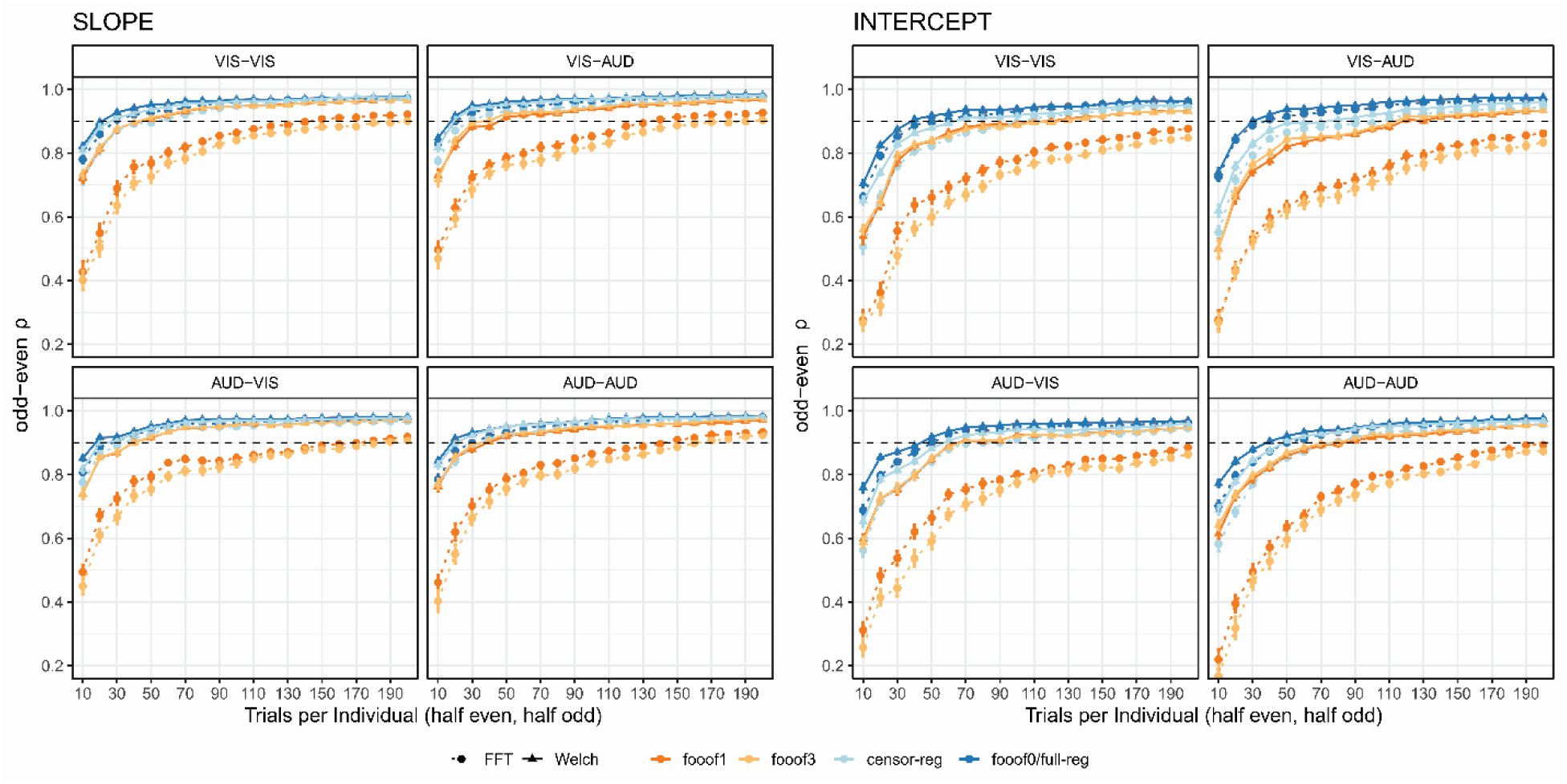
Internal consistency (odd-even reliability) as a function of trial count for slope (left) and intercept (right) across aperiodic estimation methods, shown separately for each version of the stop-signal task. The dashed horizontal line represents a reliability threshold of .90, considered acceptable. fooof0 and full-reg are displayed with the same color because their curves were virtually indistinguishable from each other. *FFT*, fast Fourier transform; *Welch*, Welch’s method; *fooof0*, *fooof1*, and *fooof3*, *fooof*-derived slope estimates with the corresponding number of modelled peaks; reg-full, full regression approach; full-reg*, full regression* approach; censor-reg*, censored regression* approach.

### 3.4 Effect size estimates within experimental contexts

Results thus far suggest that internal consistency is high for each aperiodic estimation method, but it decreases as a function of maximum peak number parameter when using *fooof*. These observations, however, provide no insight into which estimation method provides the best description of the data, i.e., how likely they are to address various research questions of interest. To examine this, we assessed which aperiodic estimation method was best able to detect expected experimental effects in the two data sets.

#### 3.4.1 Resting State

In the resting state data, we first examined the magnitude of the correlation (Spearman’s *rho*) between age and the aperiodic slope as a function of aperiodic estimation method, based on the substantial previous literature predicting spectral flattening with advancing age (e.g., Voytek et al., 2015; Dave et al., 2018; Cesnaite et al., 2023; Turner et al., 2023). The expected positive relationship between age and slope (less negative slope with increasing age, **Fig. 7A**) was found with most analytic pathways (see **Fig. 7B** for the topography of slope values in the different conditions and ***Fig. S8*** for the scalp topography of correlations with age). We observed higher *rho* values for the eyes closed (EC) compared to the eyes open (EO) condition. In EC, *fooof3* led to the highest observed effect size, followed by *fooof1* and *censored regression*. After positive slope removal, the ranking changed slightly to *fooof3*, followed by *censored regression*, then *fooof1*. For EO, most analytic pathways revealed no reliable relationship between age and slope at all. Nonetheless, *fooof1* and f*ooof3* showed the highest correlation values.

**Figure 7.**
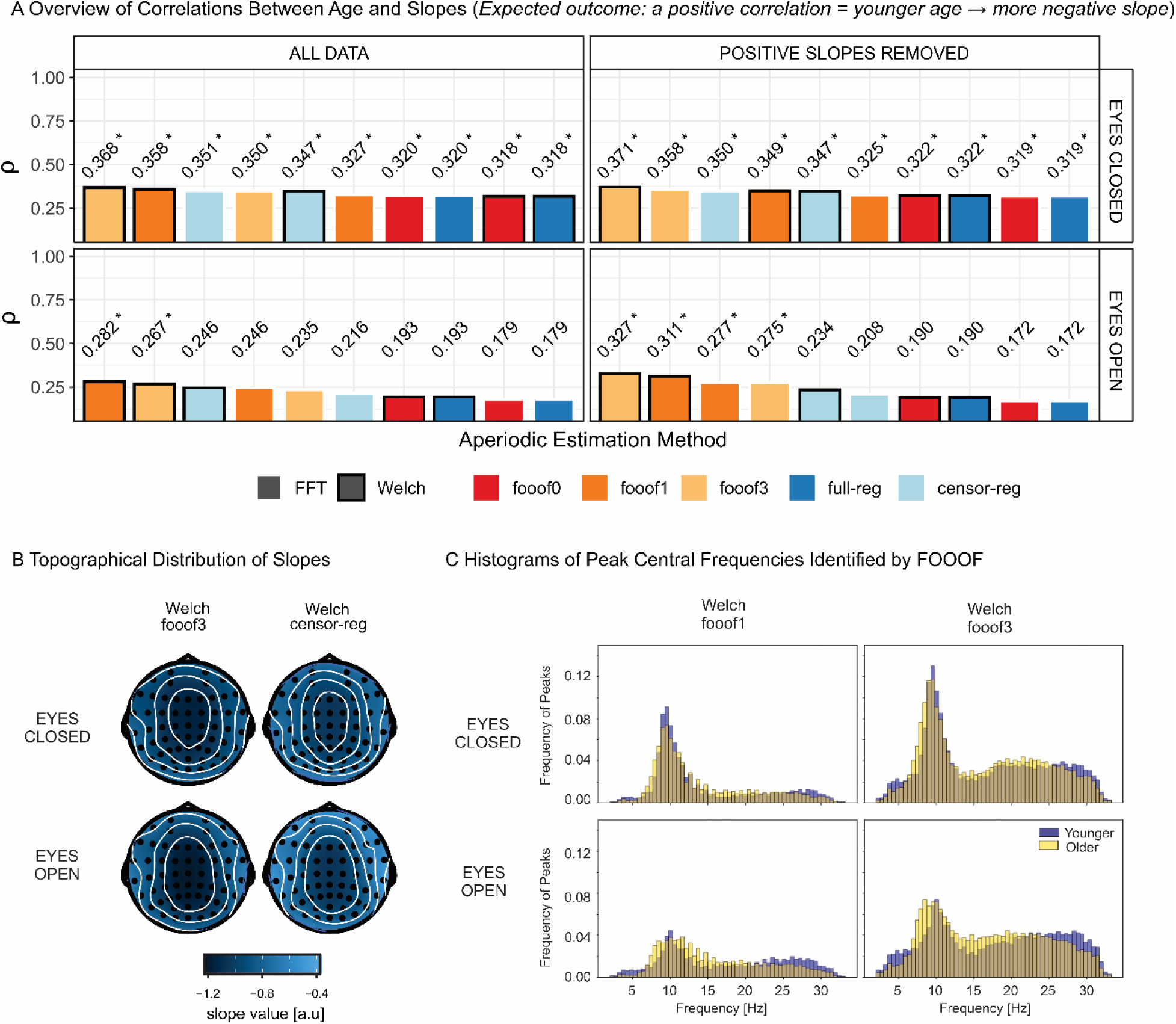
Correlations Between Age and Slope in the Resting-State Data set. (A) Spearman’s *rho* correlations between age and slope across aperiodic estimation methods, shown for the full slope data set (left panel) and after removing positive slopes (right panel), separately for eyes-closed (top) and eyes-open (bottom) conditions. Within each subplot, bars are ordered from highest to lowest correlation. Significant correlations (*p* < .05 based on the permutation test) are marked with an asterisk next to the correlation value. (B) Example topographical distribution of slopes for eyes-closed and eyes-open resting-states, shown for two selected aperiodic estimation methods: *fooof* with 3 peaks (*fooof3*) and censored regression (*censor-reg*), both derived from *Welch* spectra. (C) Histograms of peak central frequencies for younger (purple) and older (yellow) participants, identified using *Welch* with *fooof* modelled 1 peak (*fooof1*, left panel) and 3 peaks (*fooof3*, right panel), presented separately for the eyes-closed (top) and eyes-open (bottom) conditions. *FFT*, fast Fourier transform; *Welch*, Welch’s method; *fooof0*, *fooof1*, and *fooof3*, *fooof*-derived slope estimates with the corresponding number of modelled peaks; *full-reg,* full regression approach; *censor-reg,* censored regression approach.

However, it is worth considering whether *fooof*, and especially *fooof1,* might be affected by residual alpha activity only in some subjects, but not others, leading to a conflation of age-related alpha and aperiodic effects. Alpha is a periodic phenomenon prominent during rest and affected by both age and eye status (e.g., Palva & Palva, 2007, Turner et al., 2023).

To determine whether the probability of *fooof* detecting alpha activity depended on the maximum number of peaks allowed, in **Fig. 7C** we present histograms of the central frequencies of peaks identified by *fooof1* and *fooof3* (Welch-based) for the youngest (mean age = 28.14 ± 2.46, n = 17) and oldest (mean age = 64.22 ± 6.07, n = 18) thirds of the age distribution. The histograms clearly indicate a substantial increase in the number of peaks picked up in the alpha range (8-12 Hz) when more than 1-peak is allowed to be extracted. The probabilities of detecting a peak in the 8-12 Hz range based on the sum of values across this range in the standardized histograms in **Fig. 7C** for *fooof1* vs. *fooof3* in EC for younger adults were 0.45 and 0.65, respectively. For older adults these probabilities were 0.42 and 0.65. In EO for younger adults they were 0.22 and 0.42, and for older adults they were 0.27 and 0.50, respectively.

This indicates that *fooof*1 often misses alpha activity and might extract a different peak instead (likely in the beta range), opening up the possibility that undetected but present alpha peaks may bias the results in these analyses. Even in *fooof* models with multiple peaks, alpha activity may be missed if there are other larger fluctuations in the spectrum (e.g., in the EO condition a situation that may occur in which alpha amplitude is smaller than during EC). Notably, however, *censored regression* returns relatively large effects sizes, while also unequivocally accounting for alpha activity in all conditions and showing excellent reliability at the same time.

Finally, we investigated the effect size of the EC/EO difference in the slope at rest. This was partly motivated by the clear impact of eye status on the age-related slope effect, and by the theoretical expectation (supported by the arguments and findings of Zhang et al., 2019) that EC should be associated with increased inhibition (and thus a steeper spectrum/more negative slope) compared to EO, as outlined in the Introduction. This is exactly the pattern observed here for *full* and *censored regression*, and *fooof0,* regardless of spectral estimation method and of the presence or absence of positive slopes (**Fig. 8A**). Crucially, this pattern reverses, showing more negative slopes for EO compared to EC, for *fooof1* as well as *fooof3*, albeit only after positive slope removal for the latter. The scalp topographies for the different pipelines (**Fig. 8B**) reveal distributions of slope differences consistent with an alpha effect for *fooof1 and fooof3*, but not for *censored regression*, where the conditional difference is more frontally distributed. This suggests the presence of residual alpha confounds in the former but not the latter case, especially given that the effect size decreases with the number of peaks extracted (i.e., as the probability of capturing alpha increases). In summary, the problem seems to be related to the different probability of removing the alpha peak across individuals or conditions: conditions that never eliminated the alpha peak (*fooof0* or *full regression*) did not seem to show this effect.

**Figure 8.**
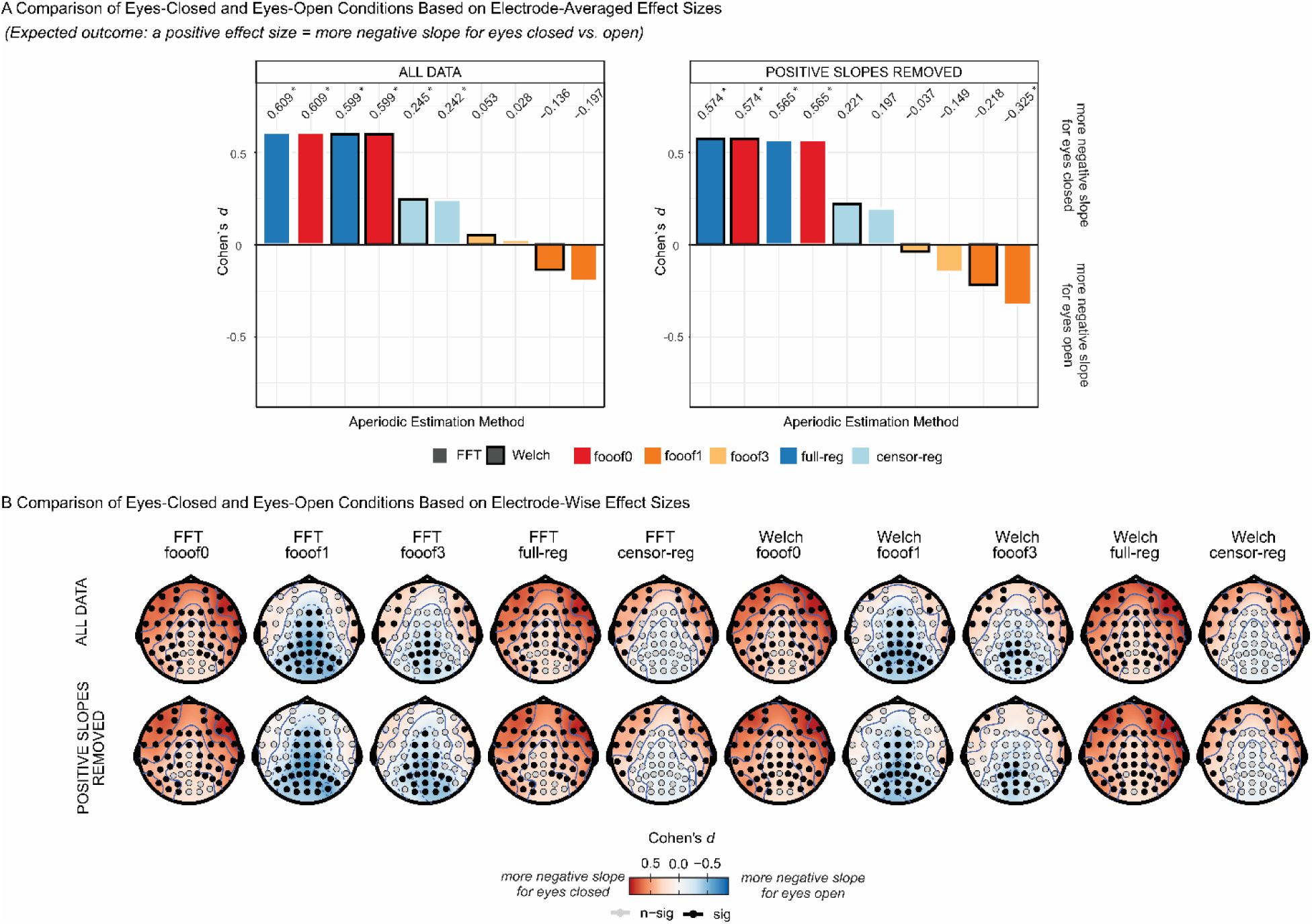
Modality Effects in the Resting-State Data set. (A) Effect sizes (Cohen’s d) for slope comparisons between eyes-closed and eyes-open resting-state, presented for the entire slope data set (left panel) and after removing positive slopes (right panel). Within each subplot, bars representing different aperiodic estimation methods are ordered from highest to lowest Cohen’s d (positive values indicate a more negative slope for eyes-closed compared to eyes-open). Significant effects (*p* < .05 based on the permutation test) are marked with an asterisk next to the d value. (B) Effect sizes (Cohen’s d) for slope comparisons between eyes-closed and eyes-open resting-state at each electrode across aperiodic estimation methods, presented for the full slope data set (top) and after removing positive slopes (bottom). Positive values indicate a more negative slope for eyes-closed relative to eyes-open. Statistical significance (*p* < .05) was assessed using a permutation test. *FFT*, fast Fourier transform; *Welch*, Welch’s method; *fooof0*, *fooof1*, and *fooof3*, *fooof*-derived slope estimates with the corresponding number of modelled peaks; *full-reg,* full regression approach; *censor-reg,* censored regression approach.

#### 3.4.2 Stop-Signal Task

In the stop-signal data set, we were interested in the difference in slope between the most vs. the least demanding conditions, as previous results suggest the former should be associated with a steeper spectrum compared to the latter, potentially reflecting the inhibition of ongoing processing (e.g., Gyurkovics et al., 2022, Kałamała et al., 2024, Lu et al., 2024). First, to demonstrate the effectiveness of the experimental manipulation, behavioral reaction times to stop-signals from this study (**Fig. 9A**) were subjected to a 2 (Target Modality) × 2 (Stop Modality) repeated-measures ANOVA. There was a significant interaction between target and stopping modalities, *F*(1,33) = 15.37, *p* < 0.001. The biggest behavioral difference was found between the VIS-VIS and the AUD-VIS conditions where the first label (AUD/VIS) refers to target modality and the second to stop modality (pairwise paired *t*-test with the largest difference: VIS-VIS vs. AUD-VIS, *t*(33) = 9.58, *p* < 0.001).

**Figure 9.**
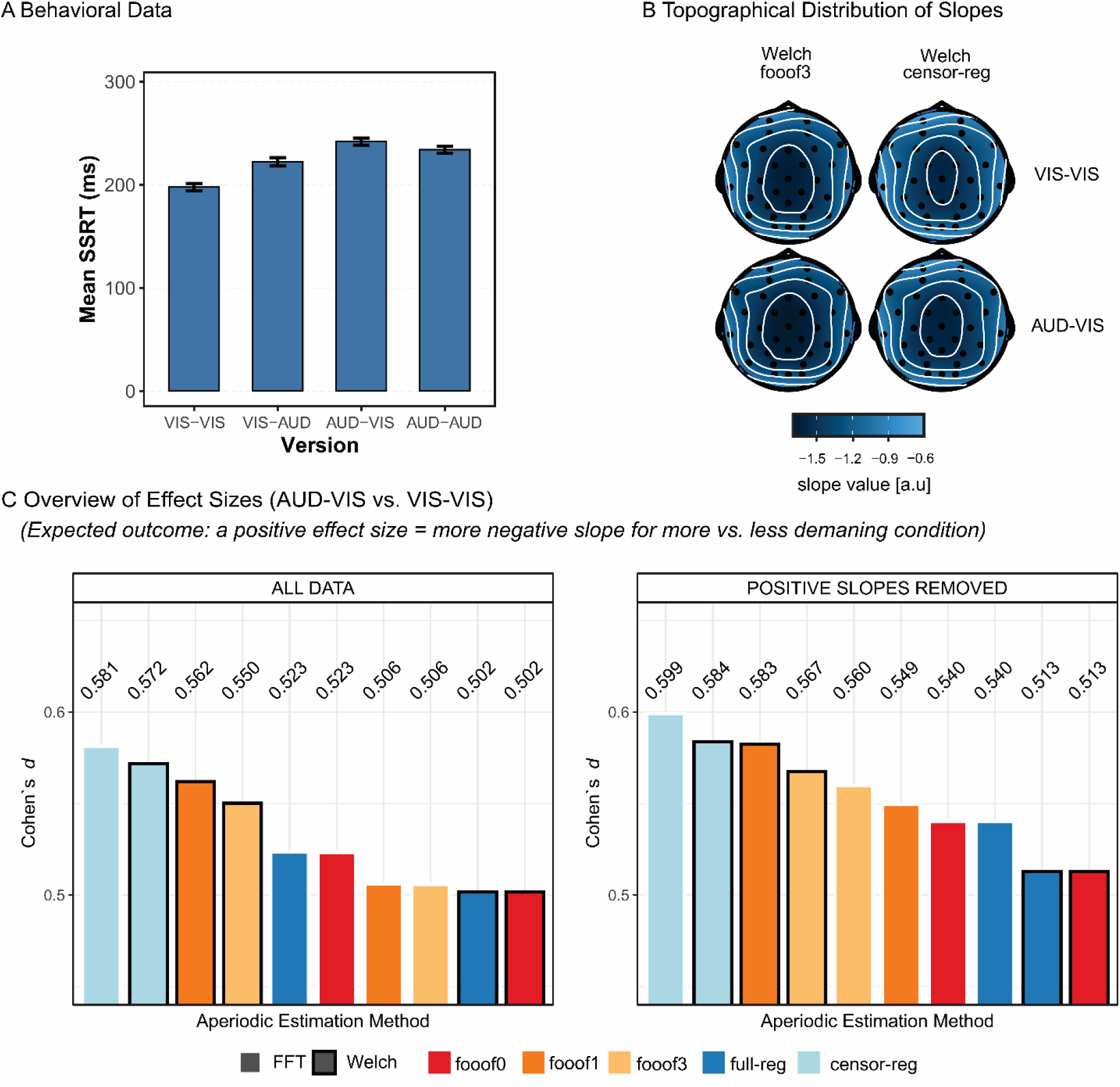
Experimental Effects in the Stop-Signal Data set. (A) Mean stop-signal reaction time (SSRT) across the four versions of the stop-signal task. (B) Exemplary topographical distribution of slopes for the least demanding (VIS-VIS) and most demanding (AUD-VIS) versions of the stop-signal task, shown for two selected aperiodic estimation methods: *fooof* with 3 peaks (*fooof3*) and censored regression (*censor-reg*), both derived from *Welch* spectra. (C) Effect sizes (Cohen’s d) for slope comparisons between AUD-VIS and VIS-VIS, presented for the entire slope data set (left panel) and after removing positive slopes (right panel). Within each subplot, bars representing different aperiodic estimation methods are ordered from highest to lowest Cohen’s d (positive values indicate a more negative slope for AUD-VIS compared to VIS-VIS). All effects are significant at *p* < .05 based on the permutation test. *FFT*, fast Fourier transform; *Welch*, Welch’s method; *fooof0*, *fooof1*, and *fooof3*, *fooof*-derived slope estimates with the corresponding number of modelled peaks; *full-reg,* full regression approach; *censor-reg,* censored regression approach.

We then contrasted the aperiodic slope estimates in these conditions and explored the effect size of the conditional difference as a function of the aperiodic estimation method to determine if the behavioral effect replicates in the EEG data, specifically in the time window preceding the “go” stimulus presentation. To this end, single-trial estimates were first averaged across epochs and then across channels. This and all subsequent analyses were conducted both before and after the removal of positive slopes. In both cases, we quantified the effect size (Cohen’s *d*) of the AUD-VIS vs. VIS-VIS difference for all aperiodic estimation methods as a measure of the pipeline’s ability to detect a theoretically meaningful effect.

As predicted, more negative slopes were found in the more demanding condition compared to the less demanding condition (see **Fig. 9B** for the scalp distribution of slope values in these two conditions). Importantly, *censored regression* yielded the largest effect sizes, both before and after positive slope removal (**Fig. 9C**, and ***Fig. S9*** in the <u>Supplementary Materials</u> for the topography of effect sizes). The extraction of multiple peaks resulted in a clear decline in effect size for these spectra. Methods that did not account for periodic activity at all (*fooof0* and *full regression*) tended to result in the smallest effect sizes.

## 4. Discussion

Because of heightened focus on aperiodic (or 1/*f*) activity in recent years in the field of cognitive neuroscience, it is of fundamental importance that its parameters are reliable and able to detect true effects. Here, we used several metrics to evaluate the adequacy of aperiodic parameter estimates, including the probability of generating artifactual results, internal consistency, and the ability to detect the hypothesized effects when they exist. Specifically, we compared different estimation methods, including the most commonly used technique, the *fooof/specparam* toolbox (Donoghue et al., 2020) with different settings (allowing 0, 1, and 3 narrowband peaks), as well as similar approaches also based on regression applied to spectra in log-log space. Across two data sets investigating different experimental issues, we found that while *fooof* generates reasonably consistent estimates, as indicated by medium-to-high odd-even reliability, its data-driven fitting procedure with several free parameters is a double-edged sword: it increases the fit of the method to actual data (which may include noise), at the cost of inflating the variability of estimates, lowering reliability and the detectability of the effects of interest. This was evident in our finding that allowing the algorithm to select progressively more peaks decreases the reliability of parameters, increases the amount of data necessary to obtain reliable estimates, and inflates the number of outlier values returned.

We also showed that a simple variation of the method, which we labeled *censored regression*, whereby a fixed range of frequencies based on theory-driven expectations is removed before regressing the log of remaining power values on the log of the corresponding frequencies, can mitigate this problem for aperiodic estimates (notably, however, it does not provide periodic estimates on its own). These findings highlight that best practices for determining aperiodic parameters should include careful consideration of *fooof* parameter settings, especially those related to the maximum number of peaks.

Unexpectedly, an even simpler method, labeled *full regression*, in which *all* frequencies are considered, and the impact of narrowband periodic activity is ignored, produced equally good and in some instances better results according to some of the metrics used in the present investigation. A possible explanation is that the range of frequencies that we excluded in the *censored regression* method are relatively close to the center of the spectrum in log-log space and therefore have a small influence on the computation of the slope (capturing the steepness of the spectrum). While at a first glance these findings could be taken to indicate that, for reliable slope estimation, peak exclusion is not always necessary, our results concerning the discoverability of effects of interest discussed below, speak against this interpretation.

It is important to note that our results indicate that the regression-based methods considered in the present study (including *fooof*) provide aperiodic parameter estimates of high internal consistency under most circumstances, as supported by good-to-excellent odd-even reliability values. This is a finding of fundamental relevance to researchers interested in quantifying aperiodic activity, as most previous studies that aimed to establish the psychometric properties of aperiodic parameters considered test-retest reliability only (Levin et al., 2020; Pathania et al., 2021; Lopez et al., 2023; Popov et al., 2023; Tröndle et al., 2023; Webb et al., 2023; Li et al., 2024; McKeown et al., 2024). While this is undoubtedly important, test-retest reliability, which measures the reproducibility of a given measure over time, cannot be meaningfully captured if the measure in question is also not internally consistent at each time point of interest, because that could lead to an underestimation of the true correlation (Spearman, 1904; Streiner, 2000). Therefore, establishing internal consistency is a crucial first step before a measure can be entered into any correlational analysis, be it the calculation of test-retest reliability, or individual differences research in general.

Because researchers use a variety of methods to pre-process their data, we also investigated the impact of different preprocessing and analytic choices on internal consistency. In our resting state data set, using simple *FFT* as opposed to *Welch’s* method to obtain the power spectra led to higher reliability for both the slope and the intercept when the *full* or *censored regression* methods (or *fooof0*) were employed for parameter estimation. However, this advantage was numerically insignificant, so based on our reliability estimates we cannot clearly recommend one method over the other, at least for estimates derived from the relatively short time windows used here. Furthermore, in our task data, combining *FFT* with *fooof* models with multiple peaks (which is the most common setting of this parameter in *fooof*) led to significantly decreased reliability and a marked increase in the number of trials required to achieve high internal consistency compared to all other combinations, for both slope and intercept. This may be because *FFT*-based spectra are less smooth compared to *Welch*-based spectra and thus provide more opportunities for noise deflections to be identified as real peaks. This effect may have been more evident in the task-based data, where trial-by-trial spectra might be more variable compared to rest, due to shorter-lived task-induced or -elicited spectral changes that may or may not occur on all trials consistently (see e.g., Gyurkovics et al., 2022, Kałamala et al., 2024).

It is, however, important to note that *Welch*’s method produces smoother input spectra at the cost of increasing the number of parameters (e.g., segment length, overlap between segments) that need to be set by experimenters (for a discussion on researcher degrees of freedom see Simmons, Nelson, & Simonsohn, 2011). This may impact the ability to compare different studies in which different choices were made for these parameters. Notably, however, our results revealed only negligible differences between the global shape of the spectra created by *FFT* and *Welch*’s method for both data sets. It is, nevertheless, likely that larger differences might appear with longer data segments that would increase frequency resolution, and thus, the jaggedness of *FFT*-based spectra. This consideration could be important for resting state analyses, where epochs of interest may easily cover several minutes of data, but less so for task-based data where epoch durations are typically close to those used here (∼1-2 s).

Of most relevance to the current article, the parameter that had the clearest impact on the reliability of aperiodic estimates was the number of peaks that the *fooof* algorithm was allowed to extract. Increasing the number of extractable peaks led to a decline in consistency and an increase in the number of trials necessary to achieve acceptable consistency for both the slope and the intercept. In both data sets, allowing the extraction of 1 or more peaks decreased odd-even reliability below 0.9, with reliability declining with each additional peak allowed. This is a particularly sobering observation given that the maximum number of possible peaks chosen in our study was 3, whereas several studies using the *fooof* algorithm allow substantially more peaks (e.g., 6 or more) or even an “infinite” number of peaks to be extracted (e.g., Seymour et al., 2022; Tröndle et al., 2023; Turner et al., 2023). The likely reason for this dip in reliability is that allowing the algorithm to select a variable number of peaks (note that even in the 1-peak model, *fooof* might extract 0 or 1 peak) from one trial to the next introduces variability in aperiodic estimation across trials (as well as across electrodes, individuals, conditions, etc.). This is because as more peaks are removed, more points will contain data where a fitted value has been subtracted from the raw value. Importantly, unless the subtraction of periodic activity accounts for such activity perfectly (i.e., there is no error in fitting and pure aperiodic activity is perfectly recovered) the increase in the number of such “corrected” points will concomitantly increase noise in the estimation of aperiodic parameters. In accordance with this logic, methods using a fixed range of points (e.g., by fitting across all points for every trial, such as for *fooof0, full regression*, or *censored regression*) showed higher reliability compared to the 1- and 3-peak models.

Although reliability is important, it is not the only criterion that should be considered when selecting an analytic approach. In fact, not all analytic steps that increase reliability are justified. The observation that methods that assume there are no peaks in the data (*full regression* and *fooof0*) are more reliable than other methods does not imply that they provide an appropriate, veridical description of the data: reliability of a measure does not inform us about its validity (Schmidt, Viswesvaran, & Ones, 2000). It is well-established that the human EEG contains oscillatory activity (Berger, 1929; Buzsáki & Draguhn, 2004) which will manifest as peaks in the power spectrum. These peaks are likely to be independent from the aperiodic activity of interest (Gyurkovics et al., 2021). Thus, it stands to reason that they need to be removed before aperiodic parameters can be estimated, as they could bias estimates of both the slope and the intercept (Donoghue et al., 2020). In the present study we were unable to estimate validity directly as we did not have access to the ground truth generative processes. We did, however, examine the ability of different methods to estimate the size of the experimental effects of interest. The logic of this approach is that a higher effect size for an empirically established and/or theoretically grounded aperiodic effect could be taken to indicate that the method used provides increased statistical power. However, this is only true if the method does not introduce biases. In cases in which the previous literature is insufficient to provide a strong hypothesis, it is difficult to use this experimental effect size criterion for evaluating a spectral parameterization method. Notably, none of the experimental effects we investigated were originally detected and reported in these data sets, avoiding circularity or overfitting of noise patterns.

In short, we assessed the ability of each preprocessing and analytic pipeline to detect predicted experimental or individual difference effects in aperiodic activity in our data. For the stop-signal data set, based on previous findings (Gyurkovics et al., 2022; Kałamała et al., 2024; Lu et al., 2024), we expected to find more negative slopes (steeper spectra) in the post-stimulus period of more, relative to less, demanding trials. This was borne out by the data, but the robustness of the observed effect was dependent on the analytic approach: for both *FFT*- and *Welch*-based spectra, censored regression yielded the largest effect size, not *full regression*, *fooof0*, or even 1- and 3-peaks *fooof* models. In other words, the most internally consistent estimates (*full regression*/*fooof0*) were not the most powerful in detecting experimental differences. Instead, the *censored regression* approach provides both high internal consistency and a strong effect size by combining two elements: 1) accounting for predictable periodic peaks by excluding them from the analysis range, and 2) ensuring that aperiodic activity was fit across the *same range* in all observations. This suggests that a valid model of real aperiodic brain data must account for (remove and/or model) periodic components in some fashion. However, a theory-driven approach to defining the frequencies affected by periodic activity may be more reliable and efficient than a data-driven approach.

We also presented effect size analyses based on the resting state data set, which further support these two conclusions, but highlight that only accounting for a single peak using *fooof* might lead to inflated effect sizes when we are considering effects that could potentially be confounded by uncorrected alpha activity (e.g., the relationship between age and the shape of the spectrum, or the impact of eye closure on aperiodic activity). The probability of detecting alpha peaks was substantially diminished when only one peak was allowed to be selected (Fig. 7C). Importantly, however, allowing the extraction of multiple peaks a) still does not guarantee that alpha activity will be detected and removed if there are other larger peaks (real or noise), and b) decreases the reliability of estimates, as demonstrated by our internal consistency findings. *Censored regression*, however, did not suffer from either of these issues, as it showed excellent reliability while accounting for alpha activity in the same fashion for all spectra.

Based on these findings, we argue that a theory-driven approach to removing peaks is more psychometrically and logically tenable than a fully data-driven approach. While a fully data-driven approach to peak extraction is intuitively appealing, it may be easily compromised by susceptibility to noise (which may appear in the form of a peak) or be insensitive to real but subtle peaks in the data (i.e., false positives and false negatives, respectively, in a signal detection framework), not to mention the psychometric concern related to decreased reliability described above, which will occur even if the extraction procedure works as intended. A fully algorithmic approach also does not take advantage of the decades’ worth of findings which have demonstrated that oscillations are mostly concentrated in certain frequency ranges in the human EEG. Let us take the example of a typical human power spectrum based on M/EEG data, where power is plotted for frequencies between 1 Hz and 30 Hz. The lower value is defined by the length of the epoch on which the spectrum is based (1 Hz would correspond to a window of at least 2 seconds so that multiple cycles of the lowest frequency can be resolved), while the higher value is determined by the signal-to-noise ratio of surface recordings, which declines sharply above 30 Hz due to the presence of noise (e.g., muscle movements), and because amplitude values there tend to be so miniscule that the dynamic range of the analog-to-digital converters would not be able to resolve them. Within this reasonably typical range, most oscillatory activity will be concentrated in the theta (4-8 Hz), alpha (8-12 Hz), and beta (12-30 Hz) ranges (Gratton, 2018). More specifically, recent findings suggest that periodic activity in the human brain is contained mostly within the alpha and beta bands (Myrov et al., 2024), with alpha being quite sustained and ubiquitous across individuals (as has been known for almost a century, Berger, 1929; Palva & Palva, 2007) and beta occurring less frequently, largely in transient bursts (Lundqvist, Miller et al., 2024). Theta activity does not always show a rhythmic, oscillatory profile and it commonly only appears in surface recordings as transient short bursts encompassing one or just a few cycles after an event requiring attentional modulation (Cavanagh & Frank, 2014; Cavanagh & Shackman, 2015).

Given this profile, it is reasonable to assume that any true oscillatory activity in surface recordings would occur in a range roughly between 4 and 25-30 Hz (if we accept that theta and beta activities are oscillatory in nature), or even a narrower band between 6 and 14 Hz (if we only consider alpha as oscillatory, as was done in the *censored regression* examples used in this paper). Therefore, we propose simply removing the alpha range or a larger range encompassing theta to beta, from all M/EEG power spectra before fitting them with regression, or a *fooof0*, model. In the current paper we showed that this *censored regression* model resulted in superior reliability compared to the *fooof1* and *fooof3* models, which also attempt to account for the presence of periodic activity before aperiodic estimation. Crucially, this solution guarantees that all spectra (for every trial, electrode, and individual) contain the same number of raw, unmodified points for quantifying the aperiodic component, thereby circumventing a source of reliability deflation. One could argue that there might be true oscillatory activity above or below this range as well, but if that were the case, those peaks would be so close to the edges of the fitting range that *fooof* would likely not be sensitive to them anyway even with peak extraction allowed (Gerster et al., 2022). Furthermore, slow delta activity (below 4 Hz) requires long time windows to be detectable, and fast gamma activity (above 30 Hz) is extremely hard to observe in surface recordings due to the low-pass filtering nature of the temporal and spatial summation inherent in M/EEG recordings. Crucially, this theoretically grounded approach to the quantification of aperiodic parameters did not only lead to more reliable aperiodic estimates, it also resulted in large effect sizes for aperiodic effects hypothesized based on earlier findings (demand effect: Gyurkovics et al., 2022; Kalamala et al., 2024; Lu et al., 2024; age effect: Voytek et al., 2015; Dave et al., 2018; Cesnaite et al., 2023; Turner et al., 2023), while also successfully avoiding possible confounds from periodic (especially alpha) activity. This highlights its practical usefulness for investigations of aperiodic activity in cognitive neuroscience.

It is also worth noting that *censored regression* produced fewer physiologically improbable slope values compared to methods allowing flexible peak extraction, regardless of spectral decomposition method. In our results, we found an inflated proportion of epochs with positive slopes using *fooof* compared to *full* and *censored regression*, regardless of spectral decomposition method during rest, and for *FFT* only during the stop-signal task. Positive slope values, indicating a positive going spectrum, where power increases as a function of frequency rather than decreases, are unlikely to occur in the brain under normal physiological circumstances (e.g., Gao et al., 2017; Brake et al., 2024). They can, however, occur as a result of noise (defined broadly as non-neural activity, e.g., high-frequency noise due to muscle movements) during the recording. Crucially, we found that even after accounting for truly positive going spectra, *fooof* still yielded a disproportionate number of positive slope estimates, which suggests that these unrealistic values may be a result of the algorithm’s estimation procedure, especially when that includes peak fitting on *FFT*-based spectra, which are more likely to contain noise fluctuations compared to *Welch* spectra. This highlights the need for careful screening of all outputs for extreme and/or improbable values before statistical tests can be conducted, as such values might bias results. Furthermore, based on the positive slope findings and our results on reliability in task-based settings, we tentatively conclude that *fooof* works more robustly when combined with *Welch*-based, as opposed to *FFT*-based spectra.

The results presented here have two major limitations. First, the *censored regression* approach can only be applied to answer research questions related to aperiodic, but not periodic activity, as it does not provide estimates of oscillatory features (e.g., power, frequency, or bandwidth) in itself. One solution might be to adopt a three-step approach, similar to that used by *fooof* (e.g., Clements et al., 2021). First the aperiodic contribution to the spectrum is estimated using the *censored regression* method. Second, this component is subtracted from the spectrum. Third, the periodic activity is estimated from the residual spectrum (for example via the same method used by *fooof*, fitting multiple Gaussians). An advantage of this approach is that the residual spectrum can be calculated in original space (not log-log space) thus allowing for the use of an additive (rather than a multiplicative) model to separately estimate the periodic and aperiodic aspects of the spectrum (see Gyurkovics et al., 2021 for a discussion of the advantages of using this approach).

Second, the specific conclusions and the recommended range of frequencies to be deleted prior to regression cannot be generalized beyond surface-level M/EEG data and therefore extended to intracranial recordings (e.g., electrocorticographic or local field potential recordings). In such cases, there may not yet be sufficient agreement about which regions of the spectrum are characterized by periodic activity and therefore should be excluded from the analysis of aperiodic phenomena. One specific frequency range where dynamics might fundamentally differ between scalp-level and intracranial recordings is the gamma range (30+ Hz). In M/EEG data, for reasons outlined above (e.g., muscle artefacts, the inherent low-pass filtering of the spatial summation involved in such recordings, the dynamic range of the amplifiers used in EEG, etc.), we consider this high frequency range to mostly be reflecting noise, either physiological or external, that should be excluded from analyses, whereas, in intracranial recordings, complex neural dynamics that could be both oscillatory and non-oscillatory in nature can contribute to this broad range (Hermes et al., 2015). As such, the range to be censored in such invasive recordings must be considered carefully. Notably, to increase the generalizability of our findings across different *scalp-level* cases, we did use two highly dissimilar data sets, with many divergent features. Furthermore, the fundamental logic of our study – i.e., that parameter settings that unnecessarily increase variability between observations within the same individual will deflate reliability and possibly lead to confounds, hampering the interpretability of findings and reducing statistical power – applies to intracranial recordings too. Thus, peak extraction parameters must be carefully considered in intracranial EEG and electrocorticography studies as well.

As a broader point, it is worth emphasizing that <u>we certainly do not advocate for all studies to censor the</u> <u>exact range that was excluded in the current study</u>. In fact, the frequency region to be censored is the only user-defined parameter of this method. It should be based on previous literature and/or on investigations of the rhythmicity of the underlying time-domain signals (e.g., Karvat et al., 2024; Myrov et al., 2024) to identify what ranges contain genuine oscillatory activity. The appropriate range might depend on sample characteristics (e.g., it might differ for children or clinical samples compared to healthy young adults). However, to minimize differences in the robustness of the estimates between groups, the range should be the same for all individuals within a study (e.g., for both children and adults in developmental studies).

Finally, the conclusions and suggestions outlined in this article apply only to aperiodic estimation methods that contain adaptive peak identification, such as the widely used *fooof* toolbox (Donoghue et al., 2020), and its extensions (e.g., Barry & De Blasio, 2021; Wilson et al., 2022). There are other methods for aperiodic parameter estimation available, such as the irregular-resampling auto-spectral analysis (IRASA; Wen & Liu, 2016), which estimates the aperiodic component of the spectrum by irregularly resampling the signal and then combining the spectra of the resampled signals. A detailed comparison of the advantages and disadvantages of *fooof*-related methods and IRASA can be found in Gerster et al. (2022).

In conclusion, in this study we investigated the internal consistency, accurate effect size representation, and other performance metrics of aperiodic parameters estimated by the popular *fooof* toolbox, as well as methods that can be considered as variants of the same fundamental procedure for spectral parametrization. In both resting state and task-based data we found generally high levels of internal consistency for slope and intercept values. It was, however, clear that internal consistency was dependent on seemingly trivial preprocessing and analytic decisions, such as the maximum possible number of peaks *fooof* could extract. To mitigate this issue, we show the advantages of implementing spectral parametrization using a *censored* spectra approach (through *fooof* or any related regression-based methods) derived from M/EEG recordings to achieve reliable and robust aperiodic measures, which can then facilitate individual differences research and group comparisons using these metrics.

## Acknowledgements

This work was supported by NIA grant RF1AG062666 to Gabriele Gratton and Monica Fabiani. Data collection for the stop-signal data set was funded by a Iuventus Plus grant (0486/IP3/2011/71) from the Polish Ministry of Science and Higher Education, awarded to Magdalena Senderecka at Jagiellonian University in Krakow, Poland.

## Supplementary Materials

## Supplementary Analyses 1: Model Fit using Information Criterion and Mean Squared Error

Model performance was quantified using mean squared error (MSE) and Akaike’s Information Criterion (AIC), both computed in log power space to meet the Gaussian error assumption. For each aperiodic estimation method, residuals were defined as the difference between the observed log power spectrum and the fitted aperiodic component; squared residuals therefore corresponded to the aperiodic-corrected power spectrum. These squared residuals were summed across frequencies to obtain a residual sum of squares (RSS).

The effective number of observations (n) corresponds to the number of frequency bins contributing to each model fit and therefore differs across methods in accordance with their modelling assumptions. Models that do not explicitly parameterize oscillatory peaks were evaluated either across the full frequency range (*full-reg*; n = 32) or with the alpha band (6–16 Hz) excluded to reduce contamination from oscillatory activity (*censor-reg*; n = 23). In contrast, models that explicitly model and separate periodic (oscillatory) components (*fooof0*, *fooof1*, *fooof3*) were evaluated across the full modelled frequency range (n = 32). Model complexity (k) is defined as the number of free parameters estimated by each approach. For models that do not explicitly parameterize oscillatory peaks (i.e., *full-reg* and *censor-reg*), k = 2 corresponds to an intercept and a slope. For *fooof* models, k reflects both aperiodic and oscillatory components. The aperiodic component contributes two parameters (offset/intercept and exponent/slope). Each oscillatory peak is described by three parameters: center frequency, amplitude, and bandwidth. Consequently, the *fooof* model without peaks (*fooof0*) has k = 2 (aperiodic parameters only), the *fooof* model with one peak (*fooof1*) has k = 5 (2 aperiodic + 3 peak parameters), and the *fooof* model with three peaks (*fooof3*) has k = 11 (2 aperiodic + 3 × 3 peak parameters).

Channel-level MSE was computed as: RSS/n, and AIC was calculated as: AIC = n · log(RSS/n) + 2k. AIC values were first computed at the channel level, then averaged within participants, and finally averaged across participants, yielding one group-level MSE and AIC estimate per experimental condition and estimation method.

Importantly, this analysis was performed only for spectra derived from the Fast Fourier Transform (FFT). Welch’s method was not included because it involves segmenting the signal into overlapping windows and averaging their periodograms, which reduces the number of statistically independent frequency estimates. As a result, the effective sample size (n) cannot be defined in a straightforward fashion as the number of frequency bins, violating the assumptions underlying the AIC and MSE formulations used here.

The AIC-based comparison evaluates the *efficiency* of different aperiodic estimation approaches, defined as the <u>trade-off between goodness of fit and model complexity, within the frequency domain that each method explicitly targets by design</u>. Specifically, data-blind models (*full-reg* and *censor-reg*) are evaluated with respect to the frequency bins used to estimate the aperiodic slope (either the full frequency range or a reduced range with the alpha band excluded), whereas data-informed (*fooof0, fooof1, fooof3*) models are evaluated with respect to the fully modelled spectrum, including frequencies that may contain oscillatory structure. As a result, AIC comparisons are most directly interpretable within families of methods that share the same modelling goal and frequency coverage: comparisons among *fooof* models differing in the number of peaks, or comparisons between full and censored regression models that explicitly test the impact of excluding oscillatory frequencies. Comparisons across method families are informative at a more general level, indicating differences in efficiency given each method’s modelling assumptions, but *should not* be interpreted as reflecting *absolute superiority* in explaining *identical* spectral data.

Accordingly, this analysis allows conclusions about whether added model flexibility, such as explicitly modelling oscillatory peaks, improves the efficiency (“fit”) of aperiodic parameterization. It does not allow conclusions about the physiological reality of oscillations, the absolute accuracy of slope estimates, or which model best explains the entire power spectrum under identical constraints. In particular, a lower AIC indicates a more parsimonious and generalizable description of the spectrum given the model’s assumptions, not a *truer* representation of neural activity.

**Figure S1.**
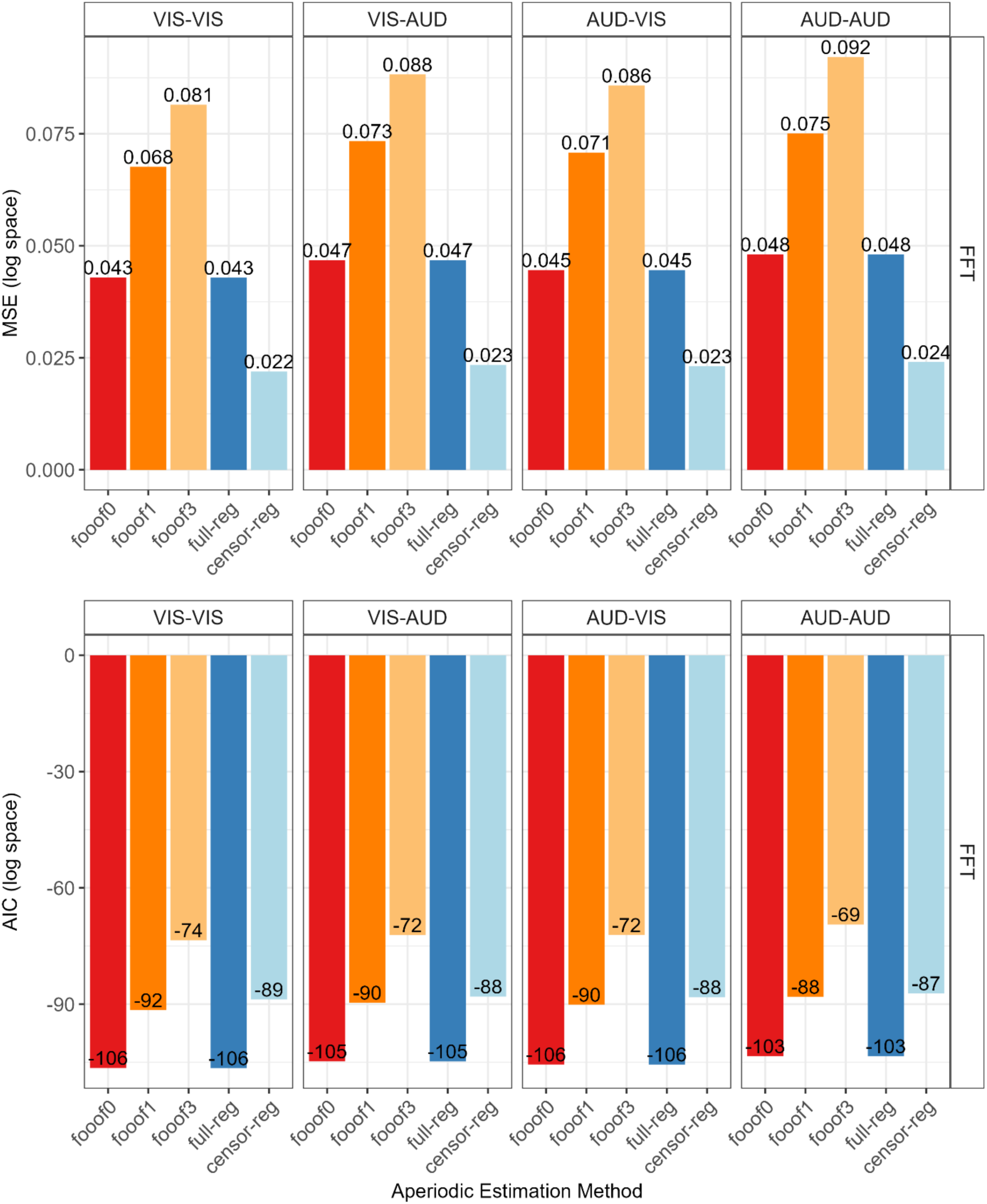
Comparison of Model-Fit Indices across Aperiodic Estimation Methods for the Stop-Signal Data set. The upper panel shows mean squared error (MSE) in log space, and the lower panel shows Akaike’s Information Criterion (AIC) in log space, for each aperiodic estimation method (*fooof0, fooof1,* and *fooof3*, *fooof*-derived slope estimates with the corresponding number of modelled peaks; *full-reg,* full regression approach*; censor-reg,* censored regression approach). Bars represent aggregated values within each condition (for details, see the text). All values were computed from power spectra derived using the Fast Fourier Transform (FFT). Lower/more negative values indicate better model fit.

Across all four task versions in the stop signal data set (Figure S1), eyes open/closed in the resting state data set (Figure S2), and both spectral estimation approaches, model comparison based on AIC consistently favored simpler aperiodic parameterizations, with the *fooof0* and the *full-reg* models yielding the same and lowest AIC values. This indicates that both approaches provide a comparably efficient balance between model fit and complexity when evaluated across the full frequency range. Importantly, increasing the number of modelled peaks did not improve model efficiency: *fooof* models with one or three peaks showed higher MSE and were strongly penalized by AIC, demonstrating that added peak parameters did not yield sufficient gains in fit to justify their complexity. Censored regression model achieved the lowest MSE, indicating superior local approximation of the aperiodic background within non-oscillatory frequency ranges, and may therefore represent a practical choice when the primary goal is accurate slope estimation rather than full-spectrum efficiency.

**Figure S2.**
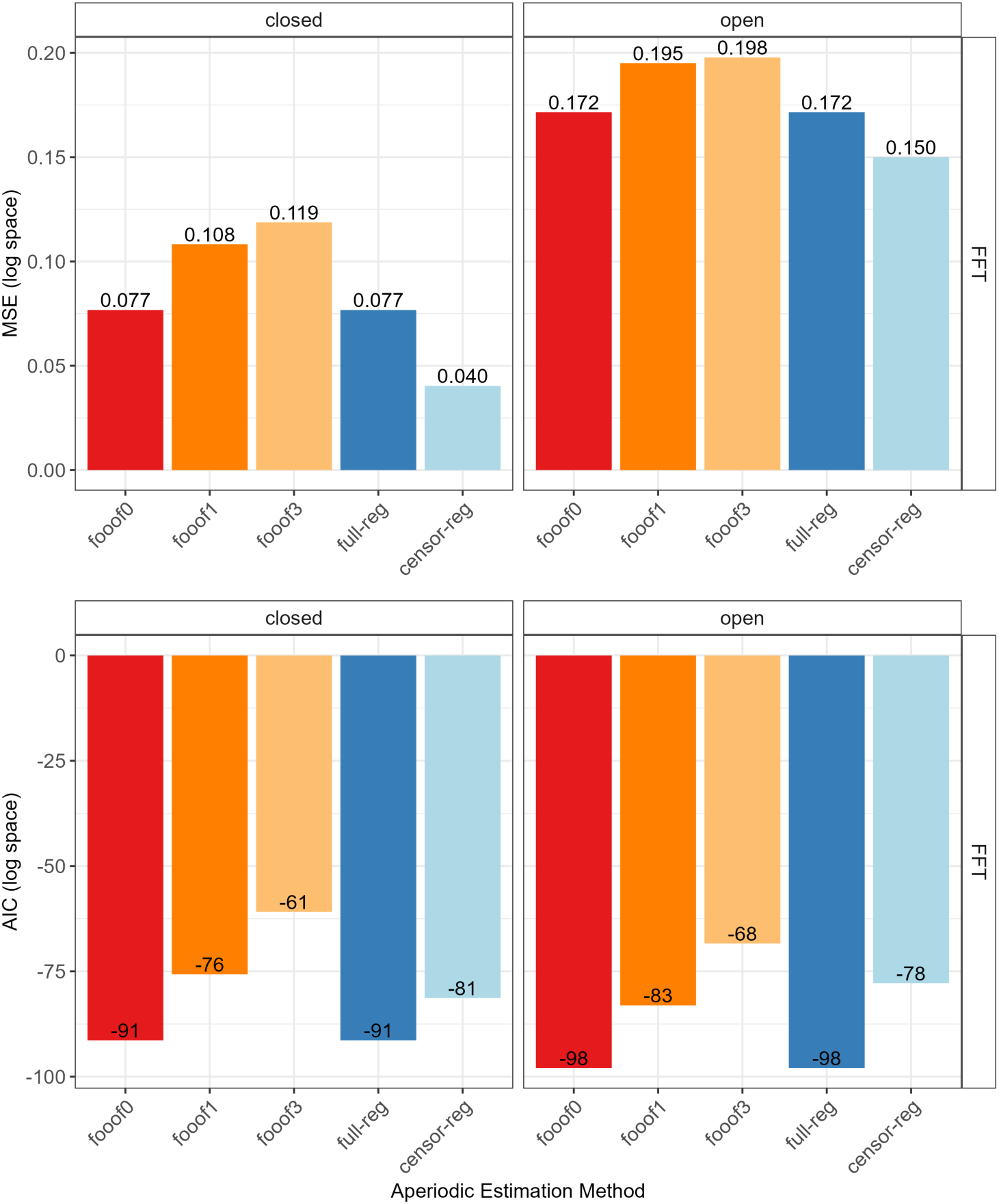
Comparison of Model-Fit Indices across Aperiodic Estimation Methods for the Resting State Data set. The upper panel shows mean squared error (MSE) in log space, and the lower panel shows Akaike’s Information Criterion (AIC) in log space, for each aperiodic estimation method (*fooof0*, *fooof1*, and *fooof3*, *fooof*-derived slope estimates with the corresponding number of modelled peaks; *full-reg*, full regression approach; *censor-reg*, censored regression approach). Bars represent aggregated values within each condition (for details, see the text). All values were computed from power spectra derived using the Fast Fourier Transform (FFT). Lower/more negative values indicate better model fit.

## Supplementary Analyses 2: Robustness of Censored Regression to Censoring Width

To evaluate the robustness of the censored regression approach to variations in the width of the censored frequency band, we conducted a simulation designed to quantify potential information loss due to censoring. We simulated 500 power spectra in log–log space as linear functions of frequency (2–33 Hz), with a ground-truth slope of −1.2 and an intercept of 1.0. To mimic realistic spectral structure, we added a Gaussian-shaped oscillatory peak centered at 10 Hz to each spectrum, along with random Gaussian noise to approximate measurement variability. For each simulated spectrum, the aperiodic slope was estimated using four approaches: (1) full regression across the entire 2–33 Hz range (“full”); (2) censored regression excluding 6–16 Hz, matching the procedure in the main manuscript (“truncated”); (3) censored regression excluding a wider frequency range than (2), i.e., 4–20 Hz (“aggressive”); (4) extreme censoring, using only the lowest and highest frequency points (i.e., two-point regression; “bookend”). Figure S3 presents the outcomes of this analysis.

**Figure S3.**
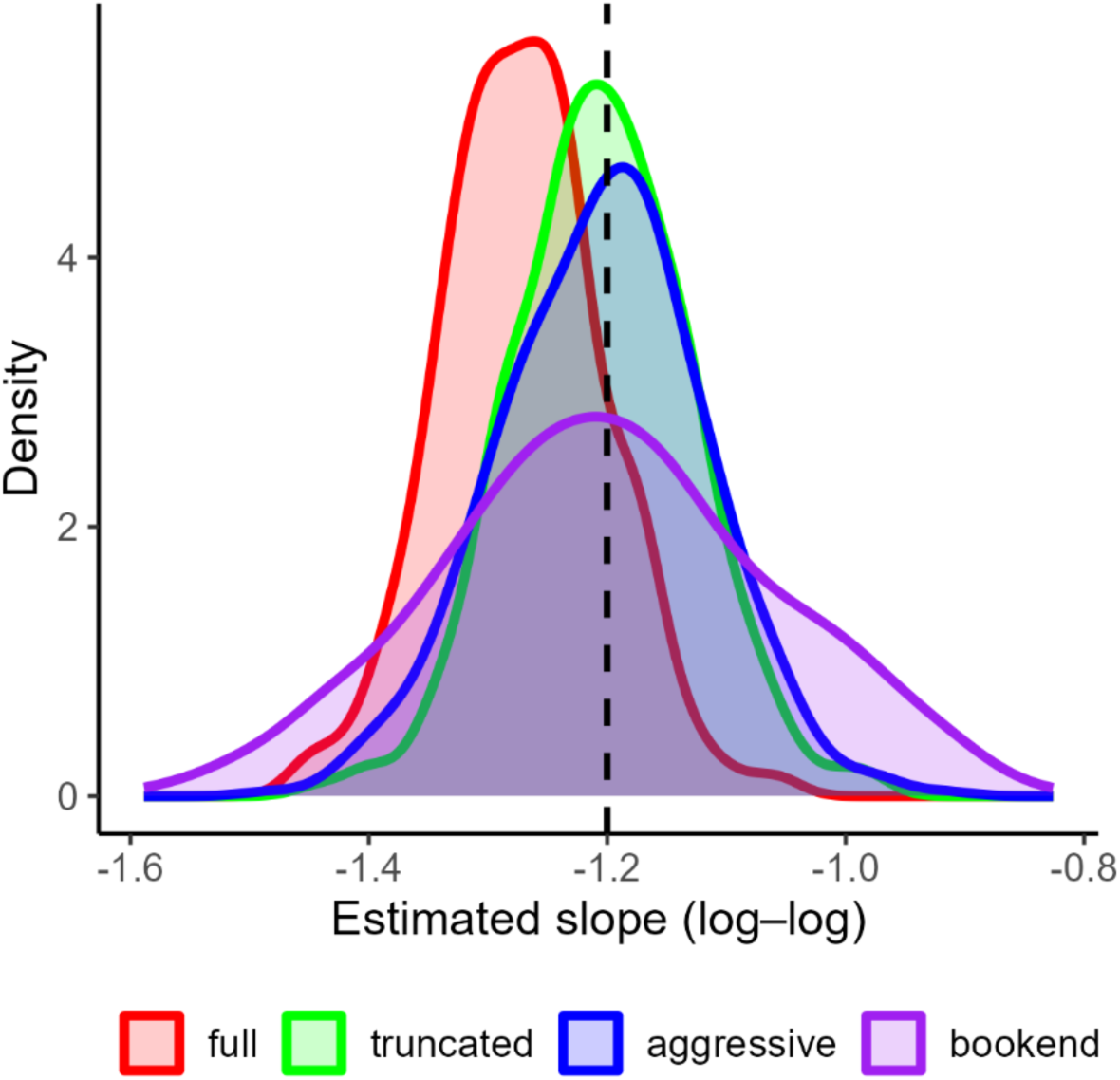
Robustness of Censored Regression to Censoring Width. Slope estimates were obtained using full regression (“full”, red), standard censoring excluding 6–16 Hz (“truncated”, green), aggressive censoring excluding 4–20 Hz (“aggressive”; blue), and extreme two-point regression (“bookend”; purple). The dashed vertical line indicates the true slope. For details, see the text.

The simulation demonstrates three key findings. First, full regression systematically misestimates the ground-truth slope, corroborating the main manuscript’s conclusion that oscillatory peaks bias conventional regression approaches when the full frequency range is used. Second, both the standard (green) and aggressive (blue) censoring procedures yield virtually identical slope estimates, despite differing substantially in the number of excluded frequency bins. Importantly, the aggressive approach removes many frequency points that are not directly contaminated by the oscillatory peak, yet it produces estimates indistinguishable from the standard censoring procedure. This indicates that moderate over-censoring results in negligible loss of information for estimating the aperiodic slope. Finally, even extreme censoring (purple; based on only two frequency points) produces slope estimates centered around the true value. However, this approach shows markedly greater variability, reflecting the increased sensitivity to noise when slope estimation relies on minimal data.

Together, these results demonstrate that censored regression is robust to reasonable variations in censoring width and that excluding oscillatory bands does not meaningfully distort recovery of the underlying aperiodic parameter.

**Table S1.** Average Slope and Intercept Values across Participants and Electrodes for Each Estimation Approach after Outlier Removal.

| Spectral<br>Decomposition | Aperiodic<br>Estimation | Slope<br>Mean | Slope<br>SD | Slope<br>Min | Slope<br>Max | Intercept<br>Mean | Intercept<br>SD | Intercept<br>Min | Intercept<br>Max |
| --- | --- | --- | --- | --- | --- | --- | --- | --- | --- |
| **Resting State Data set** |  |  |  |  |  |  |  |  |  |
| **N = 61, EEG Electrodes = 59. mean epochs = 45.45** |  |  |  |  |  |  |  |  |  |
| FFT | fooof0 | -1.06 | 0.53 | -5.19 | 0.00 | 0.89 | 0.75 | -1.39 | 6.81 |
| FFT | fooof1 | -1.11 | 0.66 | -6.37 | 0.00 | 0.71 | 0.82 | -1.89 | 7.11 |
| FFT | fooof3 | -1.20 | 0.65 | -6.38 | 0.00 | 0.69 | 0.83 | -2.10 | 7.14 |
| FFT | reg-full | -1.06 | 0.53 | -5.9 | 0.00 | 0.89 | 0.75 | -1.39 | 6.81 |
| FFT | reg-censor | -1.00 | 0.51 | -5.06 | 0.00 | 0.78 | 0.71 | -1.58 | 6.61 |
| Welch | fooof0 | -1.04 | 0.57 | -5.68 | 0.00 | 0.82 | 0.76 | -1.47 | 7.41 |
| Welch | fooof1 | -1.10 | 0.68 | -6.21 | 0.00 | 0.64 | 0.81 | -1.83 | 7.49 |
| Welch | fooof3 | -1.19 | 0.67 | -6.91 | 0.00 | 0.62 | 0.81 | -2.07 | 7.59 |
| Welch | reg-full | -1.05 | 0.57 | -5.36 | 0.00 | 0.82 | 0.76 | -1.47 | 7.41 |
| Welch | reg-censor | -0.98 | 0.51 | -5.06 | 0.00 | 0.71 | 0.71 | -1.58 | 7.20 |
| **Stop-Signal Task Data set** |  |  |  |  |  |  |  |  |  |
| **N = 34, EEG Electrodes = 32, mean epoch range 607-779** |  |  |  |  |  |  |  |  |  |
| FFT | fooof0 | -1.34 | 0.37 | -2.49 | -0.23 | 1.13 | 0.43 | -0.26 | 2.55 |
| FFT | fooof1 | -1.35 | 0.55 | -3.19 | -0.03 | 0.86 | 0.63 | -1.04 | 2.74 |
| FFT | fooof3 | -1.41 | 0.57 | -3.21 | -0.02 | 0.83 | 0.65 | -1.14 | 2.71 |
| FFT | reg-full | -1.34 | 0.37 | -2.50 | -0.23 | 1.13 | 0.43 | -0.26 | 2.55 |
| FFT | reg-censor | -1.28 | 0.37 | -2.43 | -0.15 | 1.01 | 0.44 | -0.50 | 2.40 |
| Welch | fooof0 | -1.39 | 0.34 | -2.49 | -0.35 | 1.34 | 0.40 | 0.10 | 2.72 |
| Welch | fooof1 | -1.37 | 0.39 | -2.62 | -0.22 | 1.14 | 0.43 | -0.28 | 2.57 |
| Welch | fooof3 | -1.42 | 0.38 | -2.63 | -0.25 | 1.14 | 0.43 | -0.29 | 2.53 |
| Welch | reg-full | -1.39 | 0.34 | -2.49 | -0.35 | 1.34 | 0.40 | 0.10 | 2.72 |
| Welch | reg-censor | -1.33 | 0.33 | -2.40 | -0.29 | 1.23 | 0.39 | -0.04 | 2.56 |

*Note.* Data were averaged across epochs, channels, and participants for each combination of Spectral Decomposition and Aperiodic Estimation. *FFT*, fast Fourier transform; *Welch*, Welch's method; *fooof0*, *fooof1*, and *fooof3*, *fooof*-derived slope estimates with the corresponding number of modelled peaks; *reg-full*, full regression approach; *reg-censor*, censored regression approach.

**Table S2.** Frequency of Improbable Values. All Positive Slopes, Bootstrap Analyses Ranked Results for Both Data sets.

| Spectral Decomposition | Aperiodic Estimation | Probability of Being the Best | Mean Rank<sup>a</sup> |
| --- | --- | --- | --- |
| Resting State Data set – Eyes Open |  |  |  |
| FFT | reg-censor | 98.26 | 1.04 |
| FFT | fooof0 | 1.76 | 2.48 |
| FFT | reg-full | 1.76 | 2.48 |
| Welch | reg-censor | 0.00 | 4.35 |
| Welch | fooof0 | 0.00 | 5.35 |
| Welch | reg-full | 0.00 | 5.35 |
| FFT | fooof1 | 0.00 | 7.28 |
| FFT | fooof3 | 0.00 | 7.71 |
| Welch | fooof3 | 0.00 | 9.00 |
| Welch | fooof1 | 0.00 | 10.00 |
| Resting State Data set – Eyes Closed |  |  |  |
| FFT | fooof0 | 100 | 1.50 |
| FFT | reg-full | 100 | 1.50 |
| FFT | reg-censor | 0.00 | 3.00 |
| Welch | fooof0 | 0.00 | 4.50 |
| Welch | reg-censor | 0.00 | 4.50 |
| Welch | reg-full | 0.00 | 6.00 |
| FFT | fooof3 | 0.00 | 7.00 |
| FFT | fooof1 | 0.00 | 8.00 |
| Welch | fooof3 | 0.00 | 9.00 |
| Welch | fooof1 | 0.00 | 10.00 |
| Stop-Signal Task Data set |  |  |  |
| Welch | fooof3 | 99.96 | 1.00 |
| Welch | reg-censor | 0.04 | 2.00 |
| Welch | fooof1 | 0.00 | 3.00 |
| Welch | fooof0 | 0.00 | 4.50 |
| Welch | reg-full | 0.00 | 4.50 |
| FFT | reg-censor | 0.00 | 6.00 |
| FFT | fooof0 | 0.00 | 7.00 |
| FFT | reg-full | 0.00 | 7.99 |
| FFT | fooof1 | 0.00 | 9.00 |
| FFT | fooof3 | 0.00 | 10.0 |

*Note.* <sup>a</sup> smaller is better.

**Table S3.**
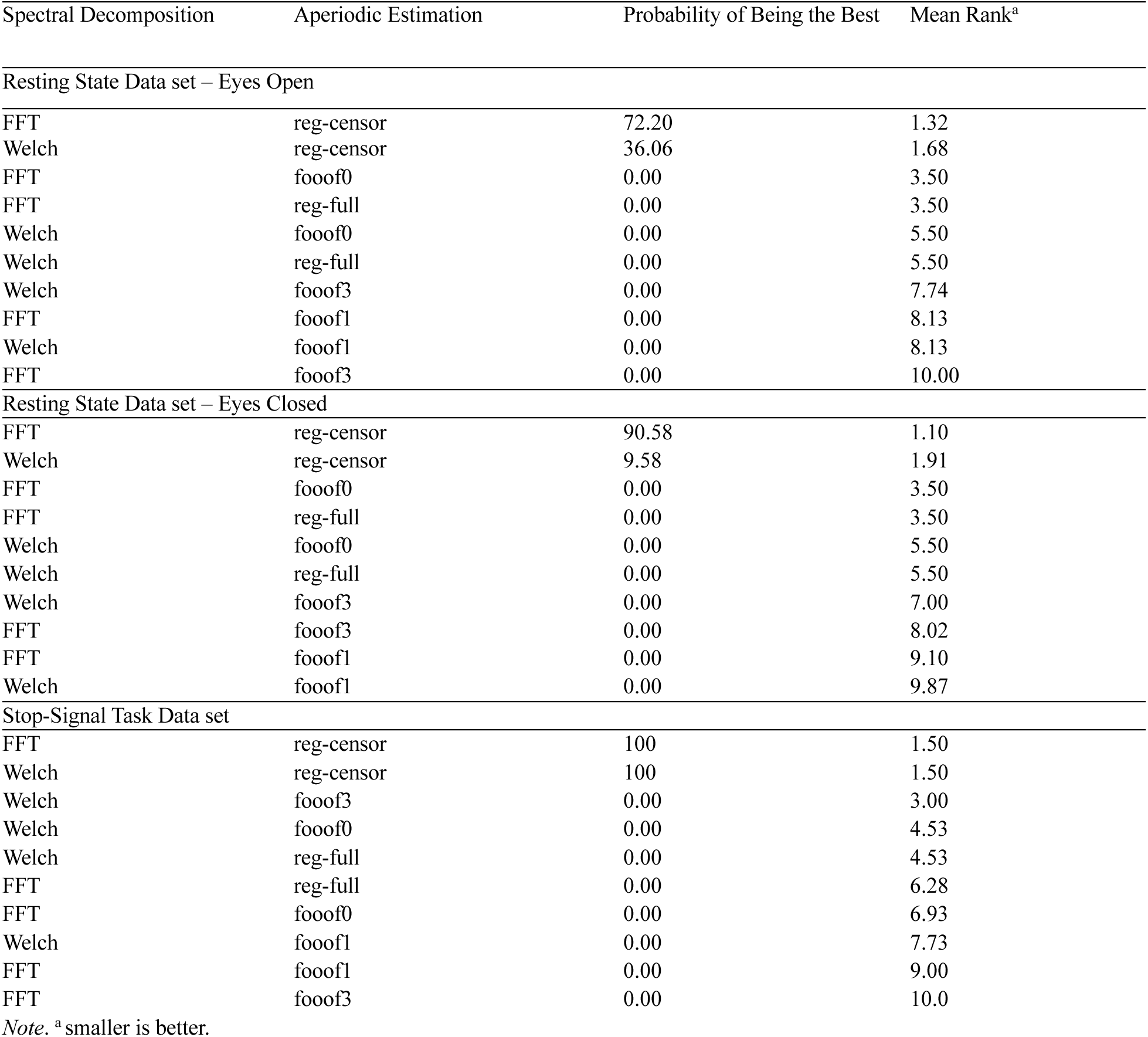
Frequency of Improbable Values. Positive Slopes Not Explained, Bootstrap Analyses Ranked Results for Both Data sets.

**Figure S4.**
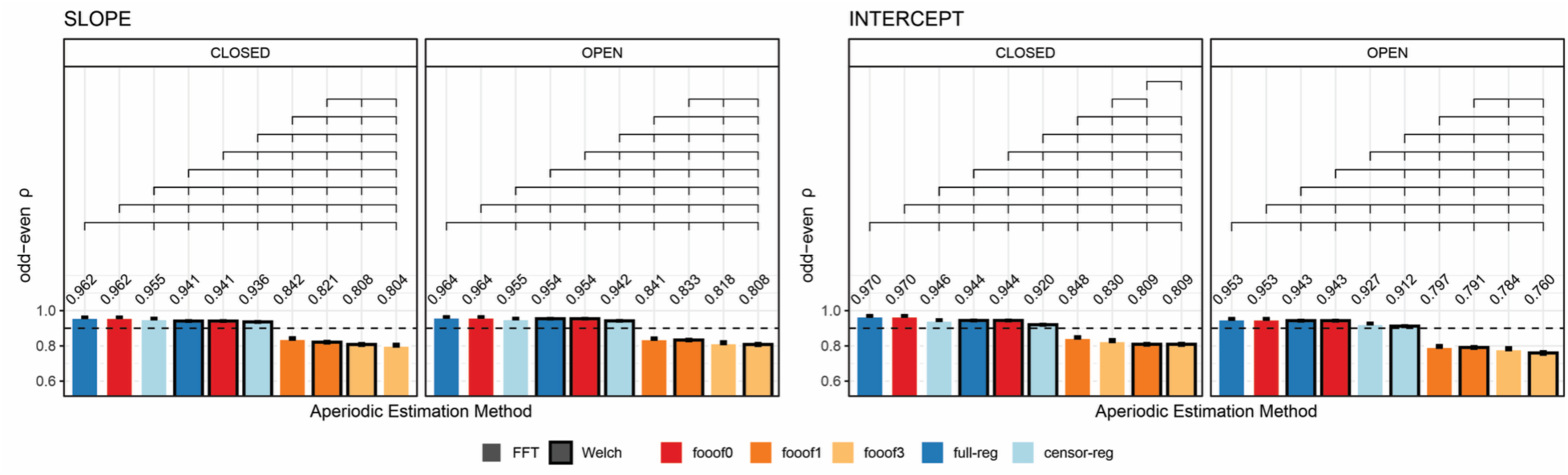
Internal consistency (odd-even reliability) after removing positive slopes for slope (left) and intercept (right) across aperiodic estimation methods, shown separately for eyes-closed and eyes-open during resting state. Within each subplot, bars are ordered from highest to lowest reliability. Brackets above the bars indicate statistically significant differences (*p* < .05 using permutation testing). Error bars reflect within-subject standard errors of the mean (SEM), computed using Morey’s (2008) correction for repeated-measures designs. The horizontal dashed line represents a reliability threshold of .90, considered acceptable.

**Table S4.**
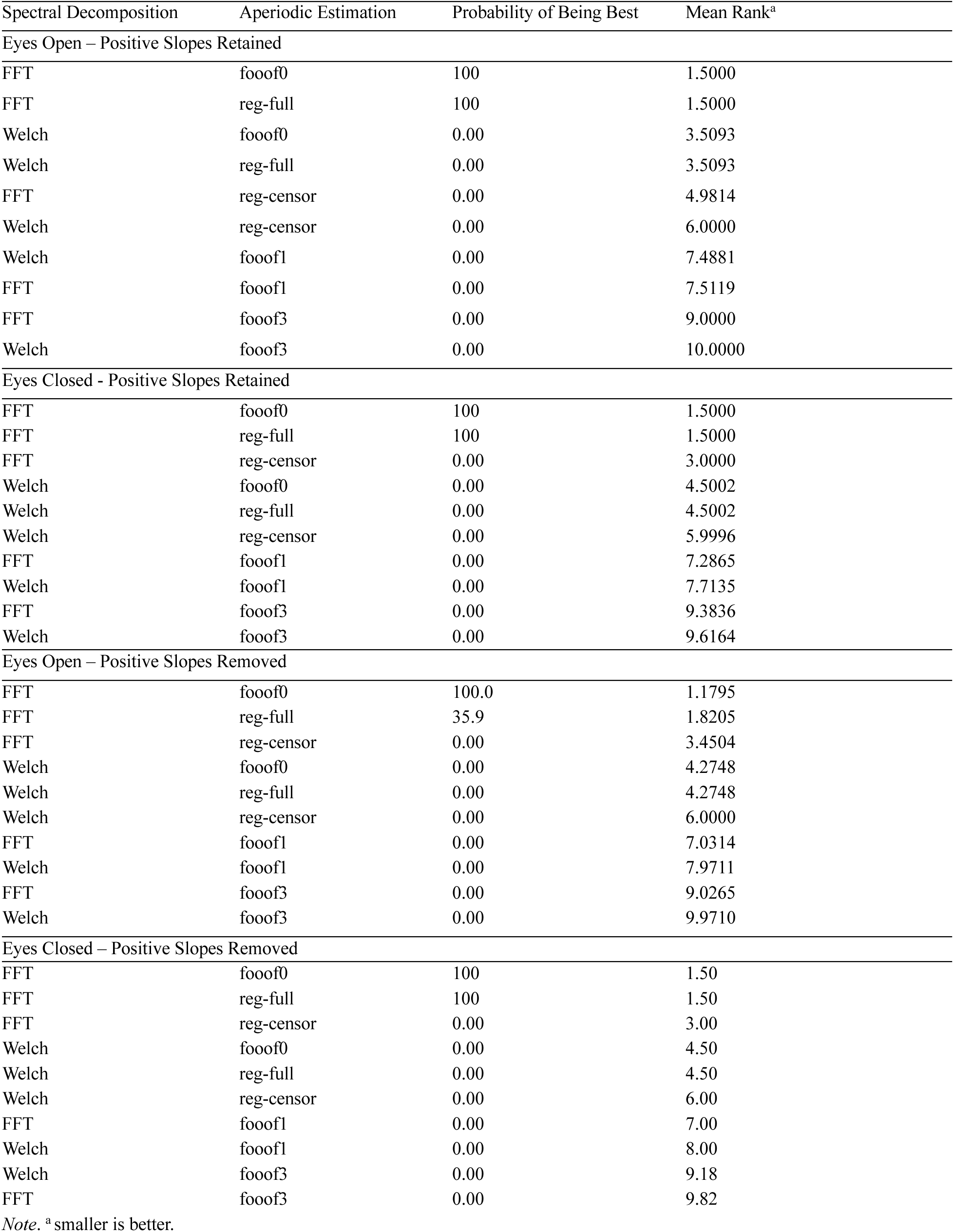
Exponent Odd-Even Reliability - Resting State Data set - Bootstrap Analyses Results.

**Table S5.** Offset Odd-Even Reliability - Resting State Data set – Bootstrap Analyses Results.

| Spectral Decomposition | Aperiodic Estimation | Probability of Being Best | Mean Rank<sup>a</sup> |
| --- | --- | --- | --- |
| Eyes Open – Positive Slopes Retained |  |  |  |
| FFT | fooof0 | 99.98 | 1.50 |
| FFT | reg-full | 99.98 | 1.50 |
| Welch | fooof0 | 0.02 | 3.50 |
| Welch | reg-full | 0.02 | 3.50 |
| FFT | reg-censor | 0.00 | 5.00 |
| Welch | reg-censor | 0.00 | 6.00 |
| Welch | fooof1 | 0.00 | 7.28 |
| FFT | fooof1 | 0.00 | 7.73 |
| FFT | fooof3 | 0.00 | 8.98 |
| Welch | fooof3 | 0.00 | 10.00 |
| Eyes Closed - Positive Slopes Retained |  |  |  |
| FFT | fooof0 | 100.00 | 1.18 |
| FFT | reg-full | 35.98 | 1.82 |
| Welch | fooof0 | 0.00 | 3.71 |
| Welch | reg-full | 0.00 | 3.71 |
| FFT | reg-censor | 0.00 | 4.58 |
| Welch | reg-censor | 0.00 | 6.00 |
| FFT | fooof1 | 0.00 | 7.01 |
| FFT | fooof3 | 0.00 | 7.99 |
| Welch | fooof1 | 0.00 | 9.30 |
| Welch | fooof3 | 0.00 | 9.70 |
| Eyes Open – Positive Slopes Removed |  |  |  |
| FFT | fooof0 | 100 | 1.50 |
| FFT | reg-full | 100 | 1.50 |
| Welch | fooof0 | 0.00 | 3.50 |
| Welch | reg-full | 0.00 | 3.50 |
| FFT | reg-censor | 0.00 | 5.00 |
| Welch | reg-censor | 0.00 | 6.00 |
| FFT | fooof1 | 0.00 | 7.12 |
| Welch | fooof1 | 0.00 | 8.00 |
| FFT | fooof3 | 0.00 | 8.87 |
| Welch | fooof3 | 0.00 | 10.00 |
| Eyes Closed– Positive Slopes Removed |  |  |  |
| FFT | fooof0 | 100 | 1.50 |
| FFT | reg-full | 100 | 1.50 |
| FFT | reg-censor | 0.00 | 3.34 |
| Welch | fooof0 | 0.00 | 4.33 |
| Welch | reg-full | 0.00 | 4.33 |
| Welch | reg-censor | 0.00 | 6.00 |
| FFT | fooof1 | 0.00 | 7.00 |
| FFT | fooof3 | 0.00 | 8.00 |
| Welch | fooof1 | 0.00 | 9.46 |
| Welch | fooof3 | 0.00 | 9.54 |

*Note.* <sup>a</sup> smaller is better.

**Figure S5.**
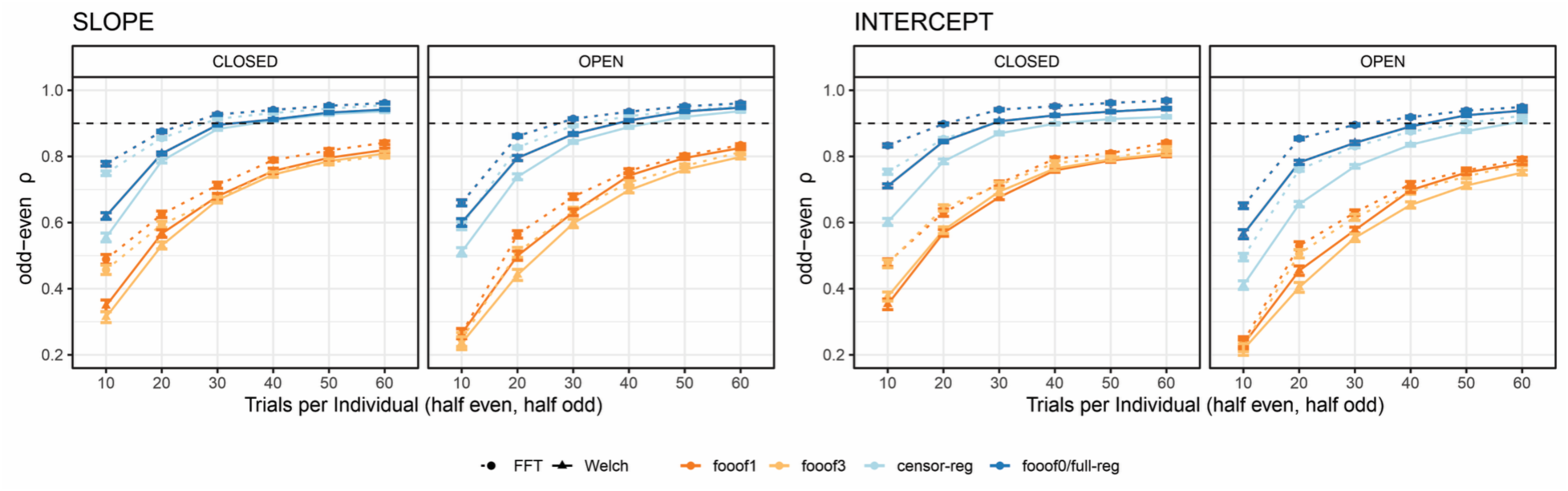
Internal consistency (odd-even reliability) as a function of trial count after removing positive slopes for slope (left) and intercept (right) across aperiodic estimation methods, shown separately for eyes-closed and eyes-open resting state. The horizontal dashed line represents a reliability threshold of .90, considered acceptable.

**Figure S6.**
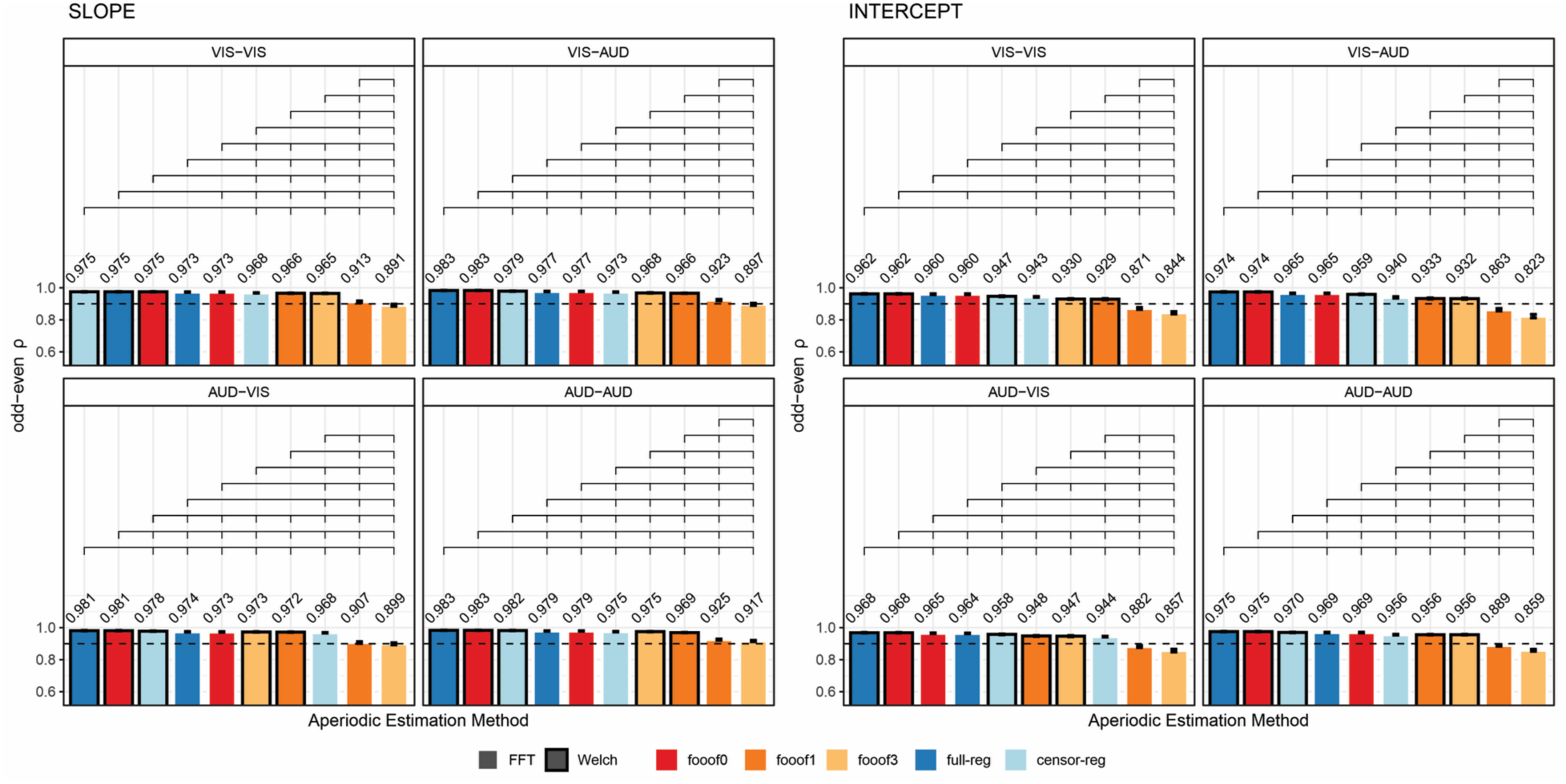
Internal consistency (odd-even reliability) after removing positive slopes for slope (left) and intercept (right) across aperiodic estimation methods, shown separately for each version of the stop-signal task. Within each subplot, bars are ordered from highest to lowest reliability. Brackets above the bars indicate statistically significant differences (*p* < .05, using permutation testing). Error bars reflect within-subject standard errors of the mean (SEM), computed using Morey’s (2008) correction for repeated-measures designs. The horizontal dashed line represents a reliability threshold of .90, considered acceptable.

**Table S6.** Exponent, Positive Slopes Retained, Odd-Even Reliability - Stop Signal Data set – Bootstrap Analyses Results.

| Spectral Decomposition | Aperiodic Estimation | Probability of Being the Best | Mean Rank<sup>a</sup> |
| --- | --- | --- | --- |
| Auditory-Auditory |  |  |  |
| Welch | fooof0 | 94.62 | 1.56 |
| Welch | reg-full | 94.62 | 1.56 |
| Welch | reg-censor | 6.12 | 2.89 |
| FFT | fooof0 | 0.00 | 4.28 |
| FFT | reg-full | 0.00 | 4.27 |
| Welch | fooof3 | 0.00 | 6.32 |
| FFT | reg-censor | 0.00 | 6.67 |
| Welch | fooof1 | 0.00 | 8.00 |
| FFT | fooof1 | 0.00 | 9.00 |
| FFT | fooof3 | 0.00 | 10.00 |
| Auditory-Visual |  |  |  |
| Welch | fooof0 | 99.98 | 1.50 |
| Welch | reg-full | 99.98 | 1.50 |
| Welch | reg-censor | 0.00 | 3.00 |
| FFT | reg-full | 0.00 | 4.80 |
| FFT | fooof0 | 0.00 | 5.24 |
| Welch | fooof3 | 0.00 | 5.67 |
| Welch | fooof1 | 0.00 | 6.30 |
| FFT | reg-censor | 0.00 | 7.99 |
| FFT | fooof1 | 0.00 | 9.00 |
| FFT | fooof3 | 0.00 | 10.00 |
| Visual-Auditory |  |  |  |
| Welch | fooof0 | 100 | 1.50 |
| Welch | reg-full | 100 | 1.50 |
| Welch | reg-censor | 0.00 | 3.07 |
| FFT | reg-full | 0.00 | 4.09 |
| FFT | fooof0 | 0.00 | 4.84 |
| FFT | reg-censor | 0.00 | 6.03 |
| Welch | fooof3 | 0.00 | 7.08 |
| Welch | fooof1 | 0.00 | 7.89 |
| FFT | fooof1 | 0.00 | 9.00 |
| FFT | fooof3 | 0.00 | 10.0 |
| Visual-Visual |  |  |  |
| Welch | fooof0 | 86.84 | 1.66 |
| Welch | reg-full | 86.84 | 1.66 |
| Welch | reg-censor | 12.82 | 2.96 |
| FFT | reg-full | 1.00 | 4.23 |
| FFT | fooof0 | 0.84 | 4.50 |
| FFT | reg-censor | 0.00 | 6.22 |
| Welch | fooof3 | 0.00 | 7.35 |
| Welch | fooof1 | 0.00 | 7.43 |
| FFT | fooof1 | 0.00 | 9.00 |
| FFT | fooof3 | 0.00 | 10.00 |

*Note.* <sup>a</sup> smaller is better.

**Table S7.** Exponent, Positive Slopes Removed, Odd-Even Reliability - Stop Signal Data set – Bootstrap Analyses Results.

| Spectral Decomposition | Aperiodic Estimation | Probability of Being the Best | Mean Rank<sup>a</sup> |
| --- | --- | --- | --- |
| Auditory-Auditory |  |  |  |
| Welch | fooof0 | 94.12 | 1.56 |
| Welch | reg-full | 94.12 | 1.56 |
| Welch | reg-censor | 6.76 | 2.88 |
| FFT | fooof0 | 0.00 | 4.30 |
| FFT | reg-full | 0.00 | 4.69 |
| Welch | fooof3 | 0.00 | 6.40 |
| FFT | reg-censor | 0.00 | 6.59 |
| Welch | fooof1 | 0.00 | 8.00 |
| FFT | fooof1 | 0.00 | 9.00 |
| FFT | fooof3 | 0.00 | 10.00 |
| Auditory-Visual |  |  |  |
| Welch | fooof0 | 99.88 | 1.50 |
| Welch | reg-full | 99.88 | 1.50 |
| Welch | reg-censor | 0.14 | 3.00 |
| FFT | reg-full | 0.00 | 4.66 |
| FFT | fooof0 | 0.00 | 5.27 |
| Welch | fooof3 | 0.00 | 5.42 |
| Welch | fooof1 | 0.00 | 6.66 |
| FFT | reg-censor | 0.00 | 7.99 |
| FFT | fooof1 | 0.00 | 9.01 |
| FFT | fooof3 | 0.00 | 9.99 |
| Visual-Auditory |  |  |  |
| Welch | fooof0 | 100 | 1.50 |
| Welch | reg-full | 100 | 1.50 |
| Welch | reg-censor | 0.00 | 3.05 |
| FFT | reg-full | 0.00 | 4.03 |
| FFT | fooof0 | 0.00 | 4.94 |
| FFT | reg-censor | 0.00 | 6.01 |
| Welch | fooof3 | 0.00 | 7.04 |
| Welch | fooof1 | 0.00 | 7.95 |
| FFT | fooof1 | 0.00 | 9.00 |
| FFT | fooof3 | 0.00 | 10.00 |
| Visual-Visual |  |  |  |
| Welch | fooof0 | 82.88 | 1.69 |
| Welch | reg-full | 82.88 | 1.69 |
| Welch | reg-censor | 18.04 | 2.74 |
| FFT | fooof0 | 0.20 | 4.34 |
| FFT | reg-full | 0.40 | 4.55 |
| FFT | reg-censor | 0.00 | 6.28 |
| Welch | fooof1 | 0.00 | 7.04 |
| Welch | fooof3 | 0.00 | 7.95 |
| FFT | fooof1 | 0.00 | 9.00 |
| FFT | fooof3 | 0.00 | 10.00 |

*Note.* <sup>a</sup> smaller is better.

**Table S8.** Offset, Positive Slopes Retained, Odd-Even Reliability - Stop Signal Data set – Bootstrap Analyses Results.

| Spectral Decomposition | Aperiodic Estimation | Probability of Being the Best | Mean Rank<sup>a</sup> |
| --- | --- | --- | --- |
| Auditory-Auditory |  |  |  |
| Welch | fooof0 | 100 | 1.50 |
| Welch | reg-full | 100 | 1.50 |
| FFT | reg-full | 0.00 | 3.58 |
| FFT | fooof0 | 0.00 | 4.12 |
| Welch | reg-censor | 0.00 | 4.30 |
| Welch | fooof3 | 0.00 | 6.93 |
| Welch | fooof1 | 0.00 | 6.96 |
| FFT | reg-censor | 0.00 | 7.12 |
| FFT | fooof1 | 0.00 | 9.00 |
| FFT | fooof3 | 0.00 | 10.00 |
| Auditory-Visual |  |  |  |
| Welch | fooof0 | 99.74 | 1.50 |
| Welch | reg-full | 99.74 | 1.50 |
| FFT | fooof0 | 0.00 | 3.07 |
| FFT | reg-full | 0.00 | 3.96 |
| Welch | reg-censor | 0.00 | 3.96 |
| FFT | fooof1 | 0.00 | 6.56 |
| Welch | fooof3 | 0.00 | 6.79 |
| Welch | reg-censor | 0.00 | 7.65 |
| FFT | fooof1 | 0.00 | 9.00 |
| FFT | fooof3 | 0.00 | 10.0 |
| Visual-Auditory |  |  |  |
| Welch | fooof0 | 100 | 1.50 |
| Welch | reg-full | 100 | 1.50 |
| FFT | fooof0 | 0.00 | 3.33 |
| FFT | reg-full | 0.00 | 3.67 |
| Welch | reg-censor | 0.00 | 5.00 |
| FFT | reg-censor | 0.00 | 6.02 |
| Welch | fooof1 | 0.00 | 7.44 |
| Welch | fooof3 | 0.00 | 7.54 |
| FFT | fooof1 | 0.00 | 9.00 |
| FFT | fooof3 | 0.00 | 10.00 |
| Visual-Visual |  |  |  |
| FFT | fooof0 | 36.50 | 2.32 |
| FFT | reg-full | 27.88 | 2.50 |
| Welch | fooof0 | 44.34 | 2.59 |
| Welch | reg-full | 44.34 | 2.59 |
| Welch | reg-censor | 0.00 | 5.14 |
| FFT | reg-censor | 0.00 | 5.86 |
| Welch | fooof1 | 0.00 | 7.19 |
| Welch | fooof3 | 0.00 | 7.81 |
| FFT | fooof1 | 0.00 | 9.00 |
| FFT | fooof3 | 0.00 | 10.00 |

*Note.* <sup>a</sup> smaller is better.

**Table S9.** Offset, Positive Slopes Removed, Odd-Even Reliability - Stop Signal Data set – Bootstrap Analyses Results.

| Spectral Decomposition | Aperiodic Estimation | Probability of Being the Best | Mean Rank<sup>a</sup> |
| --- | --- | --- | --- |
| Auditory-Auditory |  |  |  |
| Welch | fooof0 | 100 | 1.50 |
| Welch | reg-full | 100 | 1.50 |
| Welch | reg-censor | 0.00 | 3.79 |
| FFT | fooof0 | 0.00 | 3.94 |
| FFT | reg-full | 0.00 | 4.27 |
| Welch | fooof1 | 0.00 | 6.63 |
| FFT | Rec-censor | 0.00 | 7.04 |
| Welch | fooof3 | 0.00 | 7.33 |
| FFT | fooof1 | 0.00 | 9.00 |
| FFT | fooof3 | 0.00 | 10.00 |
| Auditory-Visual |  |  |  |
| Welch | fooof0 | 96.10 | 1.57 |
| Welch | reg-full | 96.10 | 1.57 |
| FFT | fooof0 | 3.98 | 2.97 |
| FFT | reg-full | 0.20 | 3.92 |
| Welch | reg-censor | 0.00 | 4.98 |
| Welch | fooof1 | 0.00 | 6.31 |
| Welch | fooof3 | 0.00 | 6.86 |
| FFT | reg-censor | 0.00 | 7.83 |
| FFT | fooof1 | 0.00 | 9.00 |
| FFT | fooof3 | 0.00 | 10.00 |
| Visual-Auditory |  |  |  |
| Welch | fooof0 | 100 | 1.50 |
| Welch | reg-full | 100 | 1.50 |
| FFT | fooof0 | 0.00 | 3.49 |
| FFT | reg-full | 0.00 | 3.51 |
| Welch | reg-censor | 0.00 | 5.00 |
| FFT | reg-censor | 0.00 | 6.14 |
| Welch | fooof1 | 0.00 | 7.35 |
| Welch | fooof3 | 0.00 | 7.51 |
| FFT | fooof1 | 0.00 | 9.00 |
| FFT | fooof3 | 0.00 | 10.00 |
| Visual-Visual |  |  |  |
| Welch | fooof0 | 87.32 | 1.73 |
| Welch | reg-full | 87.23 | 1.73 |
| FFT | reg-full | 10.62 | 2.90 |
| FFT | fooof0 | 3.74 | 3.64 |
| Welch | reg-censor | 0.00 | 5.03 |
| FFT | reg-censor | 0.00 | 5.97 |
| Welch | fooof3 | 0.00 | 7.30 |
| Welch | fooof1 | 0.00 | 7.70 |
| FFT | fooof1 | 0.00 | 9.00 |
| FFT | fooof3 | 0.00 | 10.00 |

*Note.* <sup>a</sup> smaller is better.

**Figure S7.**
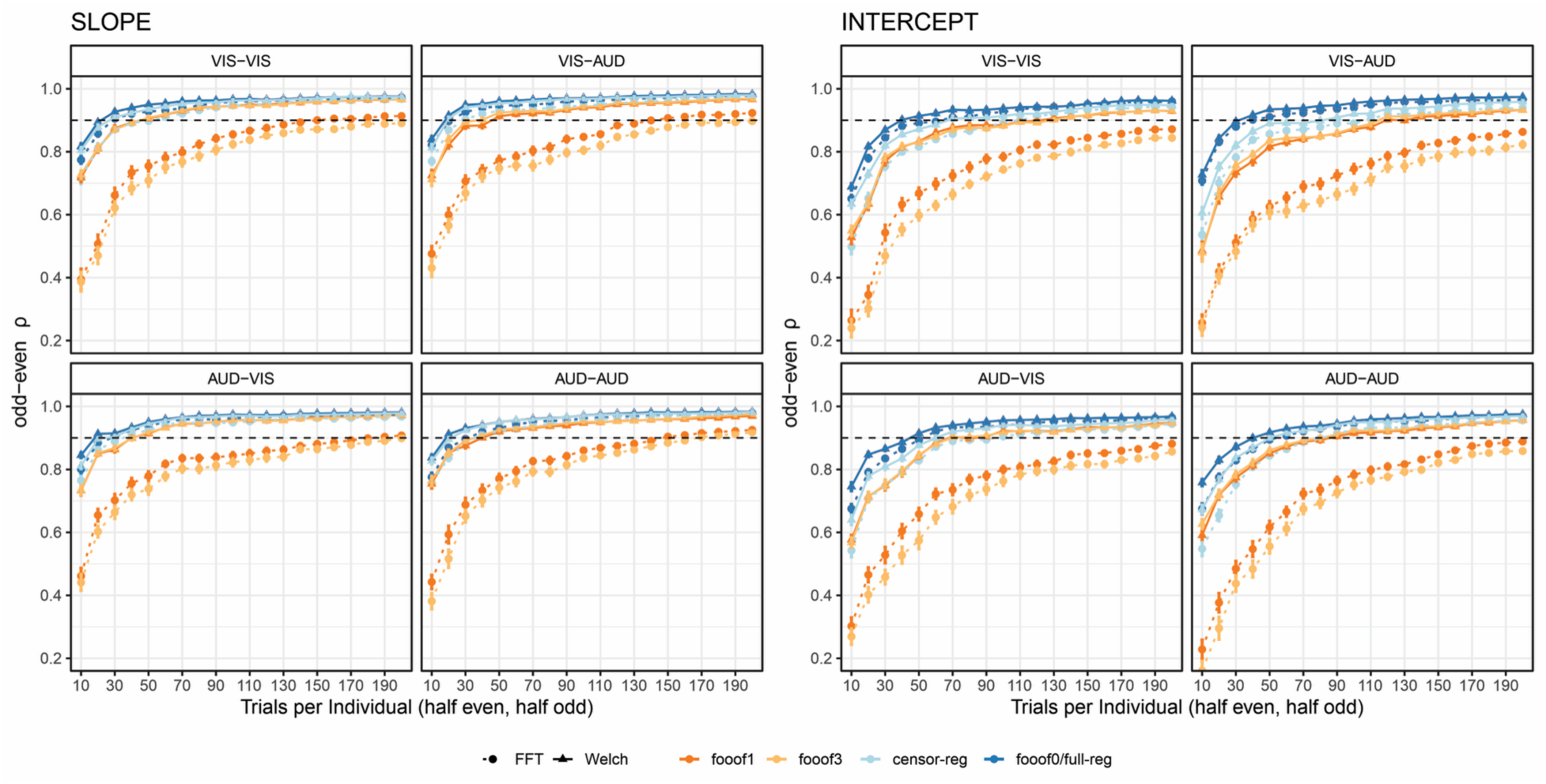
Internal consistency (odd-even reliability) as a function of trial count after removing positive slopes for slope (left) and intercept (right) across aperiodic estimation methods, shown separately for each version of the stop-signal task. The dashed horizontal line represents a reliability threshold of .90, considered acceptable.

**Figure S8.**
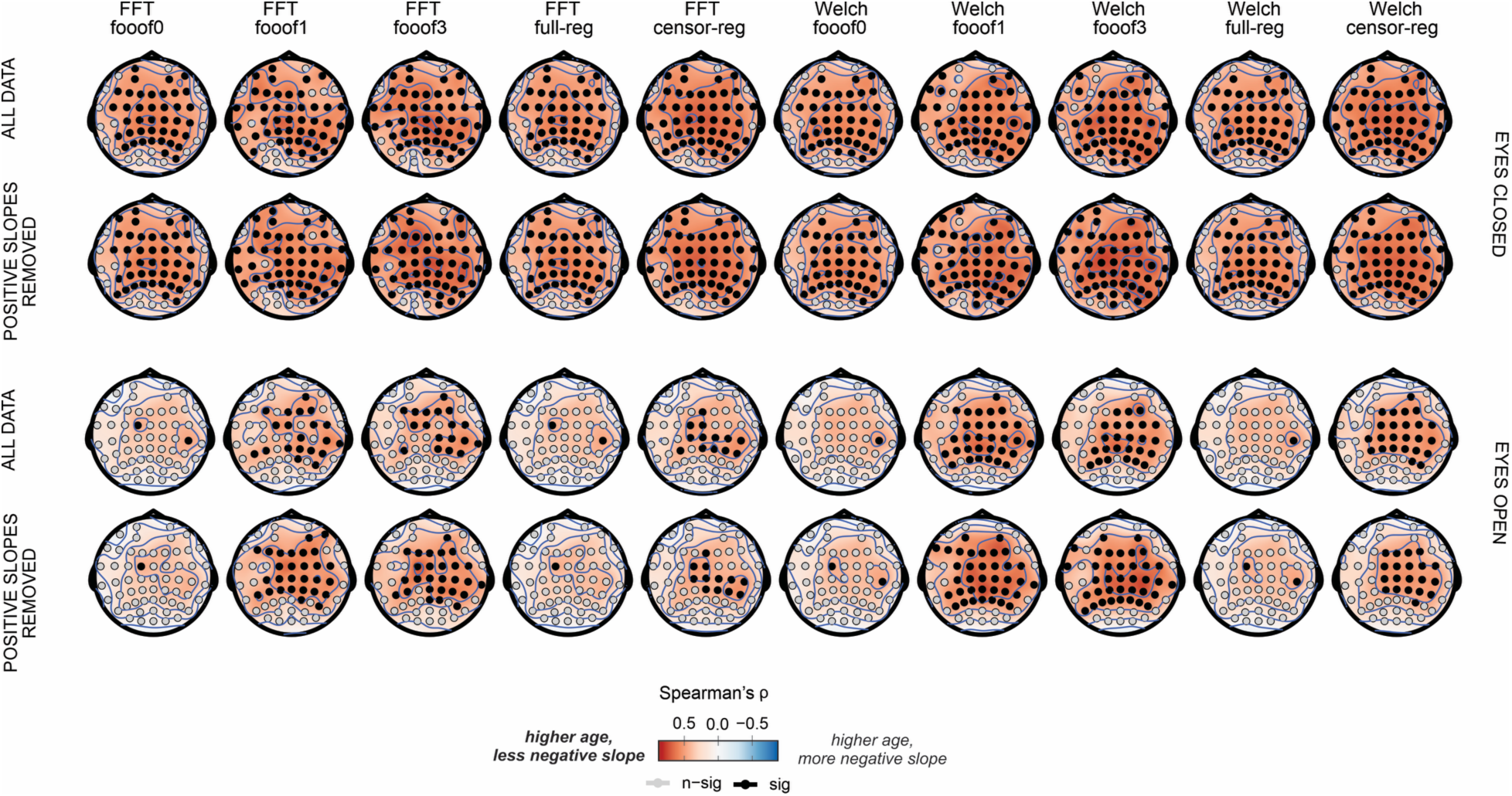
Electrode-Level Correlations Between Age and Slope in the Resting-State Data set. Spearman’s rho correlations between age and slope at each electrode across aperiodic estimation methods for eyes-closed (top) and eyes-open (bottom) resting-state conditions, shown for the full slope data set (upper) and after removing positive slopes (lower). Statistical significance (*p* < .05) was assessed using a permutation test. *FFT*, fast Fourier transform; *Welch*, Welch’s method; *fooof0, fooof1, and fooof3*, *fooof*-derived slope estimates with the corresponding number of modelled peaks; *full-reg*, full regression approach; *censor-reg*, censored regression approach.

**Figure S9.**
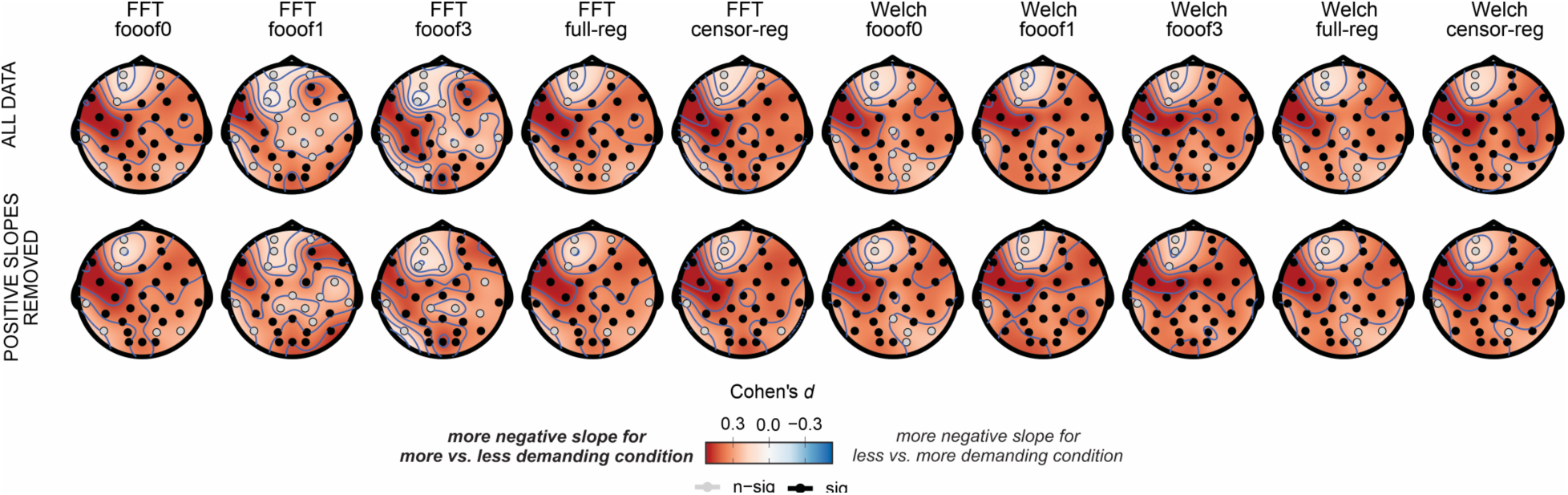
Electrode-Level Experimental Effects in the Stop-Signal Data set. Effect sizes (Cohen’s *d*) for slope comparisons between AUD-VIS and VIS-VIS at each electrode across aperiodic estimation methods, presented for the full slope data set (top) and after removing positive slopes (bottom). Positive values indicate a more negative slope for AUD-VIS relative to VIS-VIS. Statistical significance (*p* < .05) was assessed using a permutation test.

## Footnotes

1 This information can be found in the documentation materials accompanying your EEG recording system.

